# Molecular patterns and mechanisms of tumorigenesis in HPV-associated and HPV-independent sinonasal squamous cell carcinoma

**DOI:** 10.1101/2024.06.17.598514

**Authors:** Fernando T. Zamuner, Sreenivasulu Gunti, Gabriel J. Starrett, Farhoud Faraji, Tiffany Toni, Anirudh Saraswathula, Kenny Vu, Anuj Gupta, Yan Zhang, Daniel L. Faden, Michael E. Bryan, Theresa Guo, Nicholas R. Rowan, Murugappan Ramanathan, Andrew P. Lane, Carole Fakhry, Gary L. Gallia, Clint T. Allen, Lisa M. Rooper, Nyall R. London

**Author notes:** These authors contributed equally (Fernando T. Zamuner, Sreenivasulu Gunti, Gabriel J. Starrett). Nyall R. London Jr., M.D., Ph.D., F.A.C.S., Principal Investigator - Sinonasal and Skull Base Tumor Program Surgical Oncology Program (SOP), Center for Cancer Research (CCR) National Cancer Institute (NCI), National Institutes of Health (NIH) Johns Hopkins University School of Medicine Associate Professor of Otolaryngology – Head and Neck Surgery.

## Abstract

Mechanisms of tumorigenesis in sinonasal squamous cell carcinoma (SNSCC) remain poorly described due to its rare nature. A subset of SNSCC are associated with the human papillomavirus (HPV); however, it is unknown whether HPV is a driver of HPV-associated SNSCC tumorigenesis or merely a neutral bystander. We hypothesized that performing the first large high-throughput sequencing study of SNSCC would reveal molecular mechanisms of tumorigenesis driving HPV-associated and HPV-independent SNSCC and identify targetable pathways. High-throughput sequencing was performed on 64 patients with HPV-associated and HPV-independent sinonasal carcinomas. Mutation annotation, viral integration, copy number, and pathway-based analyses were performed. Analysis of HPV-associated SNSCC revealed similar mutational patterns observed in HPV-associated cervical and head and neck squamous cell carcinoma, including lack of *TP53* mutations and the presence of known hotspot mutations in PI3K and FGFR3. Further similarities included enrichment of APOBEC mutational signature, viral integration at known hotspot locations, and frequent mutations in epigenetic regulators.

HPV-associated SNSCC-specific recurrent mutations were also identified including *KMT2C*, *UBXN11*, *AP3S1*, *MT-ND4*, and *MT-ND5*. Mutations in *KMT2D* and *FGFR3* were associated with decreased overall survival. We developed the first known HPV-associated SNSCC cell line and combinatorial small molecule inhibition of YAP/TAZ and PI3K pathways synergistically inhibited tumor cell clonogenicity. In conclusion, HPV-associated SNSCC and HPV-independent SNSCC are driven by molecularly distinct mechanisms of tumorigenesis. Combinatorial blockade of YAP/TAZ and vertical inhibition of the PI3K pathway may be useful in targeting HPV-associated SNSCC whereas targeting MYC and horizontal inhibition of RAS/PI3K pathways for HPV-independent SNSCC.

**One Sentence Summary:** This study solidifies HPV as a driver of HPV-associated SNSCC tumorigenesis, identifies molecular mechanisms distinguishing HPV-associated and HPV-independent SNSCC, and elucidates YAP/TAZ and PI3K blockade as key targets for HPV-associated SNSCC.

## INTRODUCTION

Sinonasal squamous cell carcinoma (SNSCC) is an uncommon head and neck malignancy arising in the paranasal sinuses and nasal cavity. An incidence of ∼0.4 cases per 100,000 persons per year and a male predilection of ∼2:1 have been reported(*1, 2*). Smoking has been identified as a risk factor for SNSCC development although to a lesser degree than for head and neck squamous cell carcinoma (HNSCC)(*3*). Survival is poor with a 5-year survival rate of ∼50%, which may be in part due to frequent advanced stage at presentation, invasion of adjacent neurovascular structures, and other contributing factors(*1*). An improved understanding of SNSCC tumor biology is necessary to identify new potential molecular targets and to improve current therapeutic approaches.

Approximately 25% of SNSCC are associated with the human papillomavirus (HPV), compared to ∼80% of oropharyngeal squamous cell carcinomas (OPSCC)(*4, 5*). While HPV16 underlies ∼90% of HPV-associated OPSCC, HPV-associated SNSCC is associated with HPV16 in only ∼70% of cases, and other high-risk HPV subtypes are more frequently detected in SNSCC(*5–9*). HPV-associated SNSCC appears to behave differently than HPV-associated OPSCC in key aspects. Firstly, while high rates of cervical node metastasis represent a hallmark of HPV-associated OPSCC, HPV-associated SNSCC displays very low rates of cervical metastasis(*1*). Additionally, while HPV-positive tumor status confers a significant survival advantage to patients with OPSCC, this survival advantage is lower and less clear in HPV-associated SNSCC(*10*). Finally, given the rarity of SNSCC compared to OPSCC, it remains unknown whether HPV is a driver in HPV-associated SNSCC tumorigenesis or merely a bystander infection or contaminant in the sinonasal cavity(*1, 11*).

Interestingly, an additional high-risk HPV-related sinonasal carcinoma type called HPV-related multiphenotypic sinonasal carcinoma (HMSC) has been recently described(*12, 13*). This tumor is uniformly characterized by multiple histopathologic features and defined by HPV association (typically HPV33)(*13*). Although HMSC have a high local recurrence rate, these tumors rarely metastasize to the neck or distantly with only a single case of disease-related death having been reported(*13, 14*). While all cases are diffusely p16 positive, very little is known about the molecular pathogenesis of HMSC and to our knowledge no genome-wide analyses have been reported(*13*).

A variety of chromosomal aberrations have been noted in HPV-independent SNSCC including copy number variations, amplification, and microsatellite instability(*15*). Targeted sequencing has found *TP53* mutations in ∼70% of SNSCC samples(*16*). Our understanding of SNSCC tumor biology, however, remains surprisingly poor as genomics-wide analyses and high-throughput sequencing performed to date in SNSCC have been lacking. Indeed, to our knowledge only two cases of whole-exome sequencing of SNSCC have been reported as part of a larger HNSCC study(*17, 18*). Mutations in *SYNE1* and *NOTCH3* were noted in the HPV-independent SNSCC sample and, surprisingly, *TP53* was the only mutation reported in the HPV-associated SNSCC sample(*17, 18*). The rarity of SNSCC represents an obstacle to accruing a large enough cohort for comprehensive genome-wide characterization thereby limiting our knowledge of mutational frequencies and mechanistic drivers of HPV-associated and HPV-independent SNSCC and underscores a critical barrier in the advancement of novel precision therapeutics for SNSCC.

We hypothesized that performing the first comprehensive genome-wide characterization of clinically annotated cohorts of HPV-associated and HPV-independent SNSCC would reveal molecular patterns of tumorigenesis distinguishing these subtypes and define mutations linked to clinical outcomes. We also hypothesized that if HPV represented a driver in SNSCC observed mutational profiles would be shared across HPV-associated SNSCC, HNSCC and cervical cancer (CESC).

## RESULTS

### Cohort demographics and mutational analysis

The cohort consisted of 56 patients identified with squamous cell carcinoma arising from the sinonasal cavity or nasolacrimal duct. Of these 37 (66.1%) were positive on HR-HPV ISH and termed HPV-associated. P16 status for HPV-associated tumors was positive in 33 (89.2%) patients, negative in 3 (8.1%) patients, and unknown in 1 (2.7%) patient. Of HPV-independent tumors, p16 was positive in 3 (15.8%) patients, negative in 15 (78.9%) patients, and unknown in 1 (5.3%) patient. As expected, the majority of HPV-associated tumors arose in the nasal cavity while the majority of HPV-independent tumors arose in the maxillary sinus (**Table 1**)(*5*). As previously reported, patients with HPV-associated disease were younger at presentation (60.8 years) than HPV-independent (66.3 years) (**Table 1**, *P* <0.05)(*7*). Eight additional patients were identified with a diagnosis of HMSC and all were by definition positive on HR-HPV ISH. All were positive for p16 and the majority arose in the nasal cavity with a predilection for attachment sites to the inferior or middle turbinate.

**Table 1.** Patient demographics.

|  | **HPV-independent**<br>**SNSCC** | **HPV-associated**<br>**SNSCC** | **HPV-related**<br>**multiphenotypic carcinoma** |
| --- | --- | --- | --- |
| **Age at diagnosis** |  |  |  |
| Mean, years | 66.3 | 60.8 | 61.75 |
| Range, years | 40-87 | 43-75 | 45-73 |
| **Sex** |  |  |  |
| Male | 13 (68.4%) | 24 (64.9%) | 5 (62.5%) |
| Female | 6 (31.6%) | 13 (35.1%) | 3 (37.5%) |
| **Race/ethnicity** |  |  |  |
| White | 14 (73.7%) | 28 (75.7%) | 6 (75%) |
| Black | 2 (10.5%) | 7 (18.9%) | 2 (25%) |
| Other | 3 (15.8%) | 2 (5.4%) | 0 (0%) |
| **Subsite** |  |  |  |
| Nasal cavity | 5 (26.3%) | 17 (45.9%) | 7 (87.5%) |
| Ethmoid sinus | 2 (10.5%) | 5 (13.5%) | 1 (12.5%) |
| Maxillary sinus | 8 (42.1%) | 6 (16.2%) | 0 (0%) |
| Nasolacrimal duct | 3 (15.8%) | 7 (18.9%) | 0 (0%) |
| Other | 1 (5.3%) | 2 (5.4%) | 0 (0%) |
| **P16 status** |  |  |  |
| Positive | 3 (15.8%) | 33 (89.2%) | 8 (100%) |
| Negative | 15 (78.9%) | 3 (8.1%) | 0 (0%) |
| Unknown | 1 (5.3%) | 1 (2.7%) | 0 (0%) |
| **Smoking status** |  |  |  |
| Current | 3 (15.8%) | 5 (13.5%) | 1 (12.5%) |
| Former | 8 (42.1%) | 18 (48.6%) | 2 (25.0%) |
| Never | 8 (42.1%) | 14 (37.8%) | 5 (62.5%) |

Next, we performed high-throughput DNA sequencing of all HPV-independent, HPV-associated, and HMSC samples including cohorts with or without matched normal DNA. For HPV-independent SNSCC with matched normal DNA the most frequently mutated COSMIC cancer driver genes included *TP53*, *NOTCH1*, and *KRAS* (3/7 patients, 43%) as well as *CDKN2A*, *COL2A1*, *FAT4*, *FBXW7*, and *ROS1* (2/7 patients, 29%) (**Figure 1A, Supplementary Figure 1A**). Frequently mutated genes were then evaluated in our cohort of 12 additional HPV-independent SNSCC for which matched normal DNA was not available. Similar mutational frequencies were seen for *TP53*, *NOTCH1*, *CDKN2A*, *COL2A1*, and *FAT4* (**Figure 1B, Supplementary Figure 1B**).

**Figure 1.**
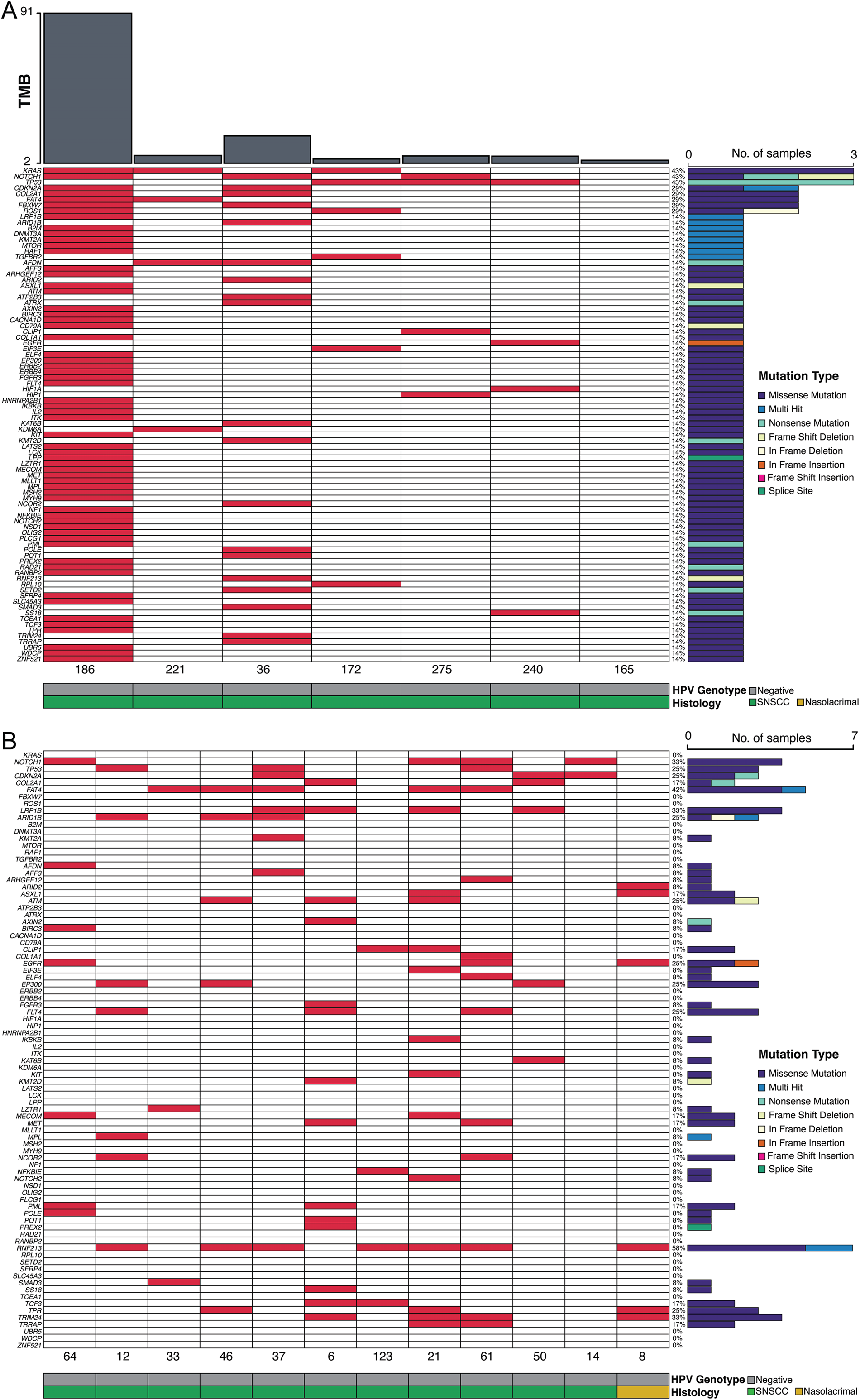
High-throughput sequencing of HPV-independent SNSCC reveals mutational patterns in COSMIC genes. **(A)** Whole-exome sequencing was performed in HPV-independent SNSCC with matched normal DNA (n=7) and somatic variants were assessed. Mutations in COSMIC genes are represented. **(B)** Whole-exome sequencing was performed in HPV-independent SNSCC without matched normal DNA (n=12) and somatic variants assessed using a panel of normal genomes. Mutations in COSMIC genes are represented.

For HPV-associated SNSCC with matched normal DNA the most frequently mutated COSMIC cancer driver genes included *KMT2D* (4/12 patients, 33%), *FGFR3* and *KMT2C* (3/12 patients, 25%), *GOLGA5*, *TET1*, and *ARID1B* (2/12 patients, 16.7%) (**Figure 2A, Supplementary Figure 2A**). Frequently mutated genes were then evaluated in our cohort of 25 additional HPV-associated SNSCC for which matched normal DNA was not available. Similar mutational frequencies were seen for *KMT2D*, *FGFR3*, *KMT2C*, and *ARID1B* (**Figure 2B, Supplementary Figure 2B**). Interestingly, recurrent in frame deletions were identified in *UBXN11* in 6/37 patients (16.2%) as well as frequent recurrent mutations in *MT-ND4*, *MT-ND5*, and *AP3S1* (**Supplementary Figure 2A-B**). Collectively, these data demonstrate a significantly different mutational profile between HPV-independent and HPV-associated SNSCC. For HMSC no recurrent COSMIC cancer driver mutations were identified in our 8-patient cohort including no patients with *KMT2D* or *FGFR3* mutations with only 1 patient with either a *KMT2C* or *ARID1B* mutation (**Figure 2A-B, Supplementary Figure 2A-B**). The tumor mutational burden (TMB) was not statistically different across the 3 groups and was relatively low with a median of 4.4 mutations per megabase (Mut/Mb) for HPV-independent, 6.6 Mut/Mb for HPV-associated, and 4.3 Mut/Mb for HMSC.

**Figure 2.**
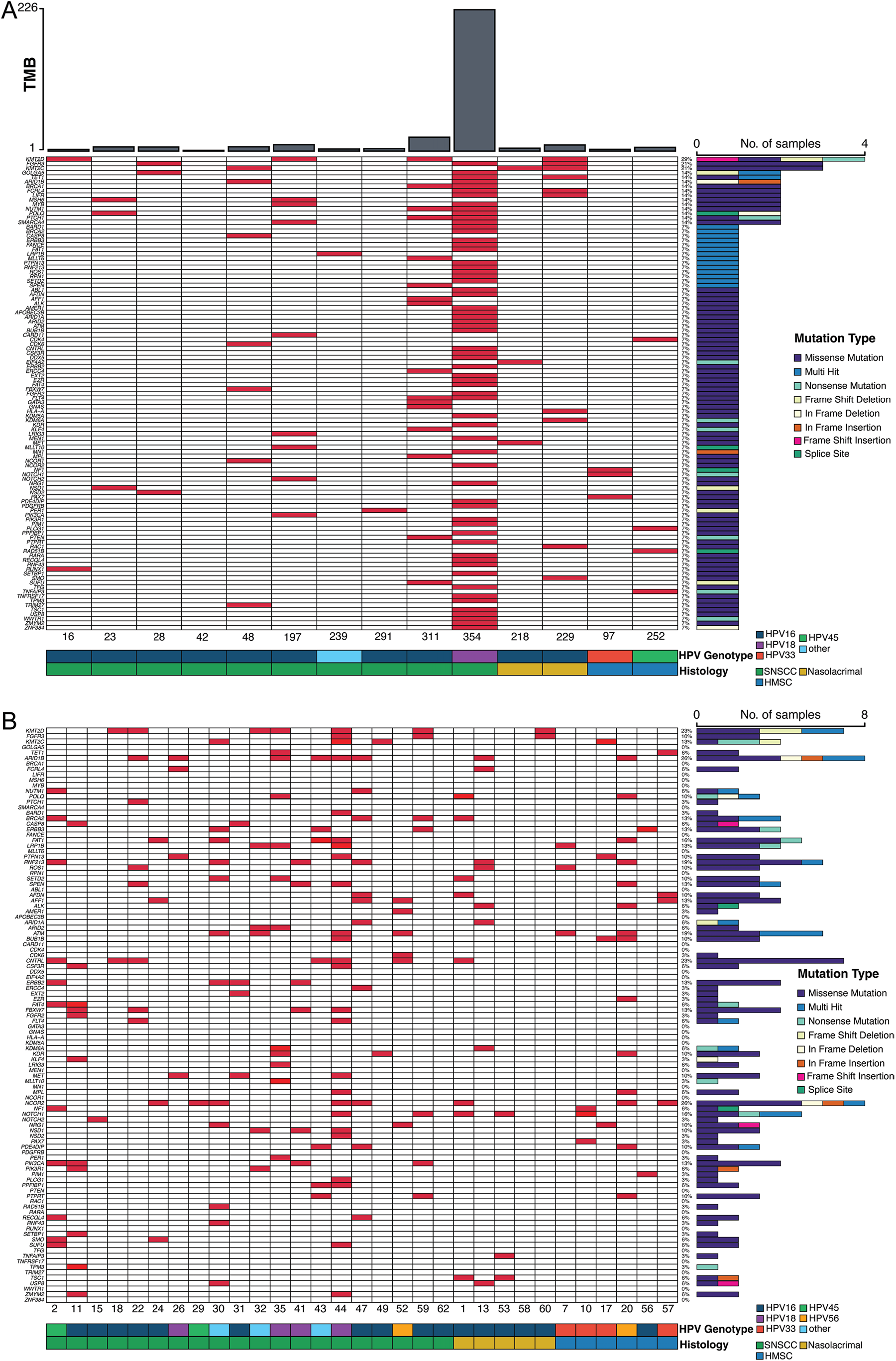
High-throughput sequencing of HPV-associated SNSCC reveals mutational patterns in COSMIC genes. **(A)** Whole-genome sequencing was performed in HPV-associated SNSCC with matched normal DNA (n=12) and HMSC (n=2) and somatic variants were assessed. Mutations in COSMIC genes are represented. **(B)** Whole-genome sequencing was performed in HPV-associated SNSCC without matched normal DNA (n=25) and HMSC (n=6) and somatic variants assessed using a panel of normal genomes. Mutations in COSMIC genes are represented.

### Identification of HPV-associated hotspot mutations

HPV-associated HNSCC and CESC have been characterized by hotspot mutations at E542K/E545K in *PIK3CA* and S249C in *FGFR3*(*19–21*). Of the 5 HPV-associated SNSCC with *PIK3CA* missense mutations 3 tumors had E542K and 1 tumor had E545K mutations (4/5, 80%) while no HPV-independent SNSCC demonstrated *PIK3CA* hotspot mutations (**Supplementary Tables 1-2**). Of the 6 HPV-associated SNSCC with *FGFR3* missense mutations 4 tumors had S249C hotspot mutations (4/6, 66.7%) while no HPV-independent SNSCC demonstrated *FGFR3* hotspot mutations. Interestingly, recurrent mutations were noted in HPV-associated SNSCC in *KMT2C* N729D (3/6, 50%), and *AP3S1* P158L (5/5, 100%) with recurrent in-frame deletions in *UBXN11* c.1464_1541 (6/8, 75%), and recurrent frameshift insertion/deletions or nonsense insertions in *MT-ND4* at c.237-238 and 4 additional individual sites (7/15, 46.7%), and in *MT-ND5* at c.891-892, c.1278-1279, and 4 additional individual sites (8/17, 47.1%) (**Figure 2A-B; Supplementary Tables 3-4**). None of these mutations were observed in HPV-independent SNSCC (**Figure 1A-B; Supplementary Tables 1-2**). Next, we evaluated the presence of SNSCC hotspot mutations in HNSCC and CESC in TCGA. Interestingly, while many *KMT2C* and *MT-ND5* mutations were noted in HNSCC or CESC, no mutations were noted at *KMT2C* N729D or *MT-ND5* c.891-892 or c.1278-1279 as seen in HPV-associated SNSCC (**Supplementary Figure 3A-D**). Mutations in *AP3S1*, *UBXN11*, and *MT-ND4* were not frequently observed or were absent in HNSCC and CSCC, and none of the recurrent changes in these genes were seen as in HPV-associated SNSCC (**Supplementary Figure 4A-F**).

Similar to HPV-associated HNSCC and CESC, mutation signature analysis revealed enrichment of APOBEC-associated mutational signature (COSMIC SB13) in HPV-associated SNSCC compared to HPV-independent SNSCC (**Figure 3A-C**; *P* < 0.05). Even though a history of smoking was present in 11/19 (57.9%) of HPV-independent SNSCC and 26/45 (57.8%) of HPV-associated SNSCC and HMSC, no enrichment in smoking mutational signature (COSMIC SBS4) was noted in any of these populations. Other signatures which did demonstrate enrichment did not demonstrate statistically significant differences when comparing HPV-associated SNSCC to HPV-independent SNSCC (**Supplementary Figure 5A-B**).

**Figure 3.**
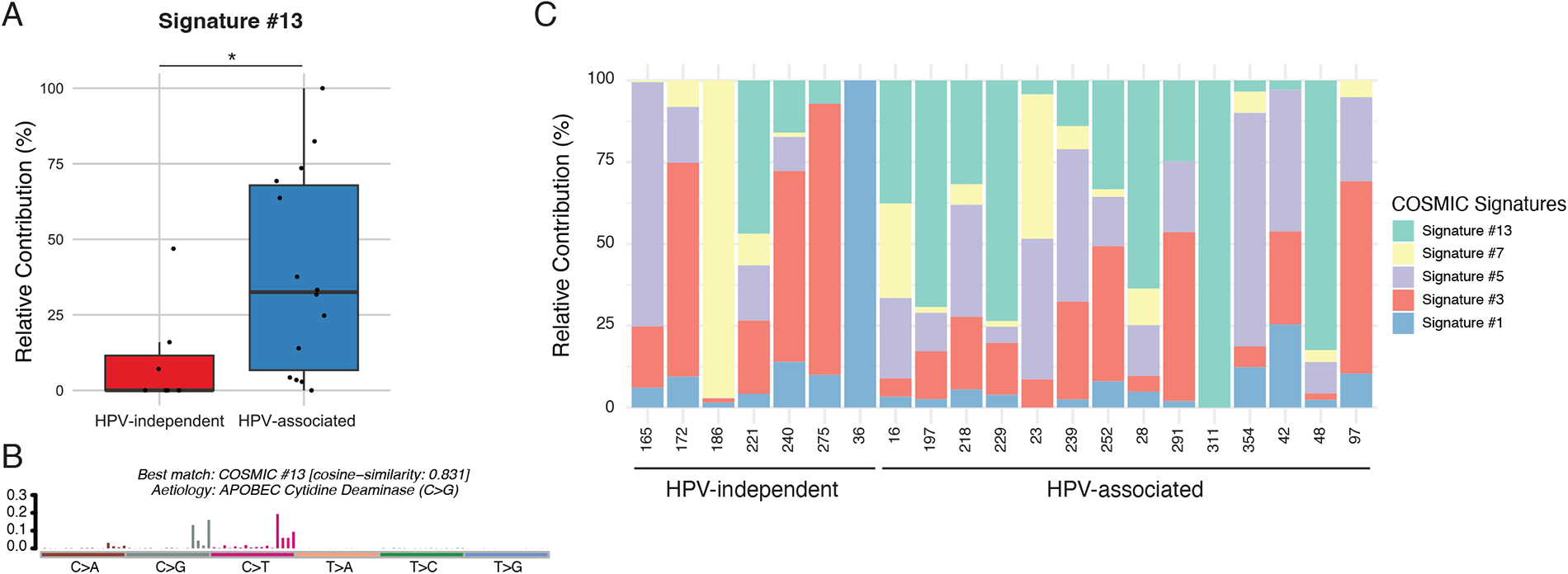
APOBEC pathway signature 13 is enriched in HPV-associated SNSCC. **(A)** APOBEC signature 13 in the HPV-independent SNSCC compared to HPV-associated SNSCC cohort with matched normal DNA. * *P* < 0.05. **(B)** APOBEC signature 13 in the HPV-associated SNSCC cohort. **(C)** Relative COSMIC signature contribution in HPV-independent and HPV-associated SNSCC.

### Correlation of mutational status with overall survival

Next, we correlated clinical outcomes with the presence of frequent mutations in HPV-independent and HPV-associated SNSCC. As previously seen for HPV-independent HNSCC(*22, 23*), the presence of a *TP53* mutation correlated significantly with worse overall survival in HPV-independent SNSCC (**Figure 4A**, *P* ˂ 0.01). No other statistically significant differences were noted with other genes of interest in HPV-independent SNSCC. Mutations in *KMT2D* and *FGFR3* in HPV-associated SNSCC were associated with worse overall survival (**Figure 4B-C**, *P* ˂ 0.05 and *P* = 0.0516, respectively) while a correlation was not seen for other genes of interest. To our knowledge, a correlation with *KMT2D* and survival has not been reported for HPV-associated HNSCC or CESC. We evaluated TCGA and found a correlation with *KMT2D* mutation and decreased overall survival in HPV-associated CESC (*P* ˂ 0.05) but not HPV-associated HNSCC (*P* = 0.412) (**Figure 4D-E**). For *FGFR3* mutations no significant difference in overall survival was noted for HPV-associated HNSCC and there were insufficient patients with *FGFR3* mutations in TCGA to evaluate for an association with CESC (**Supplementary Figure 6**).

**Figure 4.**
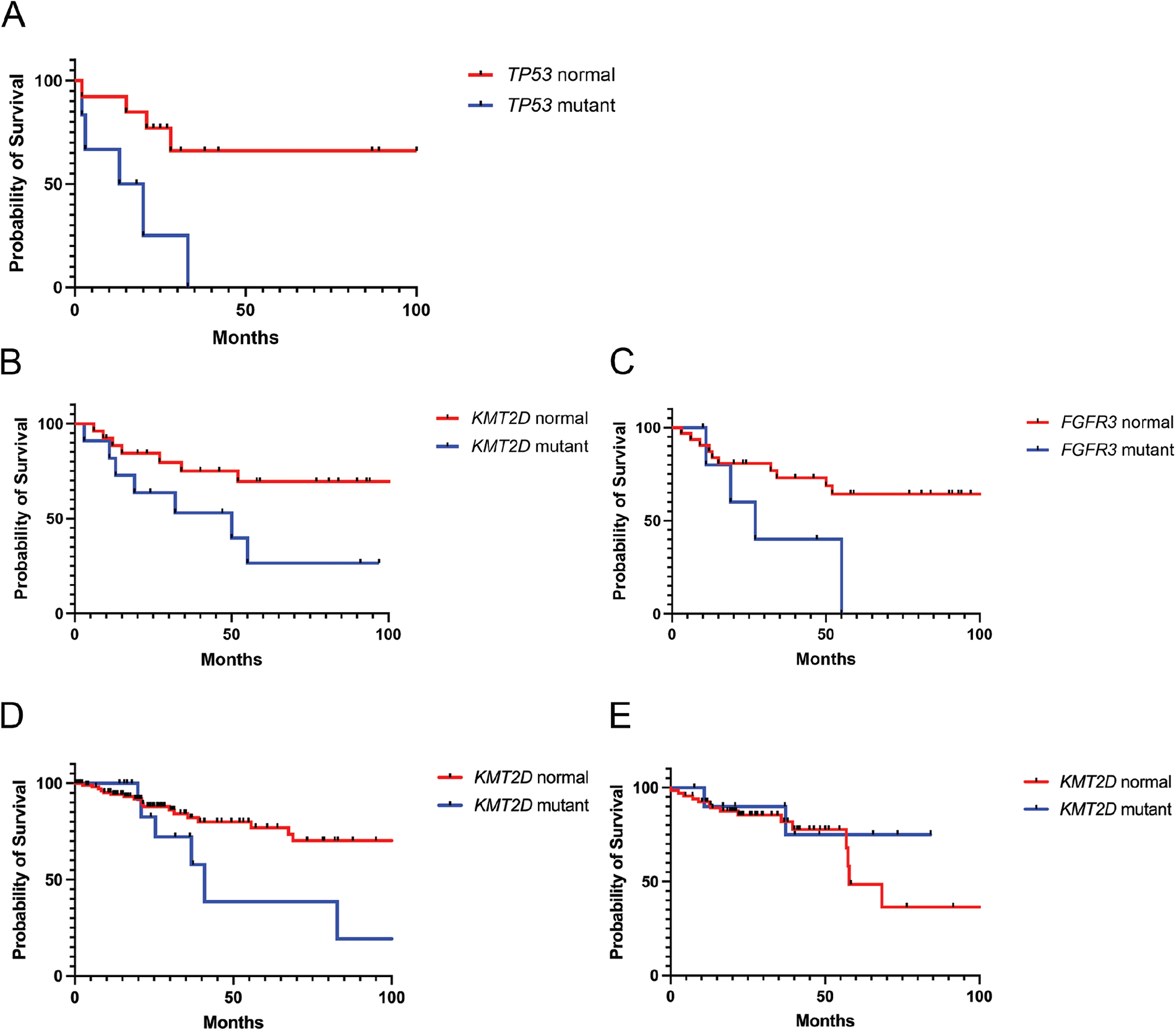
Clinical associations with overall survival and mutation status. **(A)** HPV-independent SNSCC with *TP53* mutations demonstrate an association with significantly decreased overall survival (P ˂ 0.01). **(B)** HPV-associated SNSCC with *KMT2D* mutations demonstrate an association with significantly decreased overall survival (P ˂ 0.05). **(C)** HPV-associated SNSCC with *FGFR3* mutations demonstrate an association with decreased overall survival (P = 0.0516). **(D)** HPV-associated CSCC from TCGA with *KMT2D* mutations demonstrate an association with decreased overall survival (P ˂ 0.05). **(E)** HPV-associated HNSCC from TCGA with *KMT2D* mutations and overall survival (P = 0.412).

### Assessment of viral integration

Another characteristic of HPV-associated HNSCC and CESC is frequent viral integration events in approximately 70-80% of HPV-associated tumors(*24–26*). While many of these appear random or associated with common fragile sites, several hotspots have been identified near genes involved in epithelial stem cell maintenance including *FGFR*, *TP63*, *KLF5*, *SOX2*, and *MYC*(*6, 27*). We thus hypothesized that analysis of whole genome sequencing for HPV integration events may reveal similar patterns of viral integration in HPV-associated SNSCC as seen for HPV-associated HNSCC and cervical cancer. Indeed, a wide variety of HPV integration events were identified across the genome (**Figure 5, Supplementary Table 5**). Interestingly, HMSC tended to have a higher quantity of viral integration events compared to HPV-associated SNSCC. Upon closer evaluation we noted that five tumors had integration sites near *FGFR3* (patient #7 and 56), *TP63* (patient #28 and 252), and *KLF5* (patient #60) (**Supplementary Table 5**). Interestingly although HMSC comprised a minority of our cohort, three of the five patients with integration sites near these genes had HMSC. Additionally, for patient #56 the integration appears related to a *TACC3::FGFR3* fusion as one segment of the integration event is in *TACC3*, one segment is upstream of *FGFR3*, and the final segment is at the 3’ end of *FGFR3*. Out of the 88 observed integration events on autosomes, 28 overlapped fragile sites and integrations were observed in 3 samples in *FRA2S* (sample #2, 7, and 26) and in 2 samples in *FRA1K* (sample #1, 7), *FRA6B* (sample #7, 17), and *FRA19B* (11, 311). A surprisingly high enrichment for integration events were observed in SINE, LINE, and simple repeats; however, this was strongly skewed by a few samples including #7, 11, and 218.

**Figure 5.**
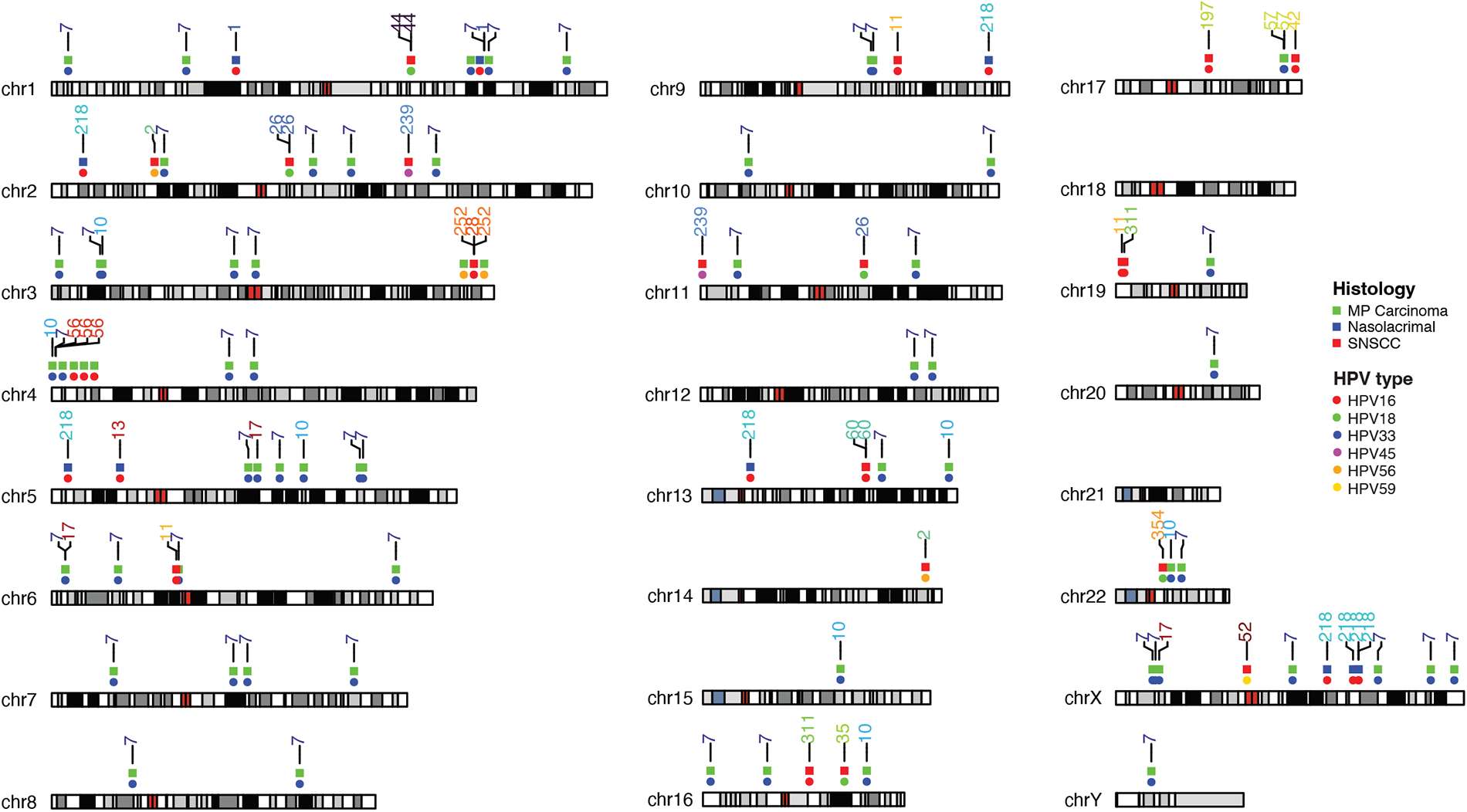
Viral integration site analysis for HPV-associated SNSCC and HMSC. Viral integration sites for HPV-associated SNSCC and HMSC are displayed for each chromosomal location. At each integration site the top row indicates the histology type while the bottom row indicates the HPV serotype.

### Elucidation of dysregulated pathways and molecular targets

Next, we sought to integrate somatic coding mutation frequencies from HPV-associated and HPV-independent SNSCC with CNV analysis to elucidate dysregulated pathways and mechanisms of tumorigenesis. Significant PI3K pathway activation was observed in HPV-associated SNSCC, driven in large part by hotspot mutations in *PI3K* and *FGFR3* in HPV-associated disease (**Figure 6**). Interestingly, HPV-associated SNSCC also demonstrated activation of the YAP/TAZ pathway (driven predominantly by copy number amplification) as well as through inactivation of FAT family protocadherin inhibitors of the Hippo pathway (**Figure 6**). The degree of YAP/TAZ pathway activation was comparible to HPV-associated HNSCC and HPV-associated CESC (**Supplementary Figure 7**). When evaluating HPV-independent SNSCC, increased pathway activity was noted through both the PI3K and RAS pathways as well as MYC (driven predominantly by copy number amplification) (**Figure 6**). Activation of RAS and MYC pathways was enhanced in HPV-independent SNSCC compared to HPV-independent HNSCC (**Supplementary Figure 8**). Collectively, these results suggest that combinatorial blockade of YAP/TAZ and vertical inhibition of the PI3K pathway may be useful in targeting HPV-associated SNSCC whereas targeting MYC and horizontal inhibition of RAS/PI3K pathways for HPV-independent SNSCC(*28*).

**Figure 6.**
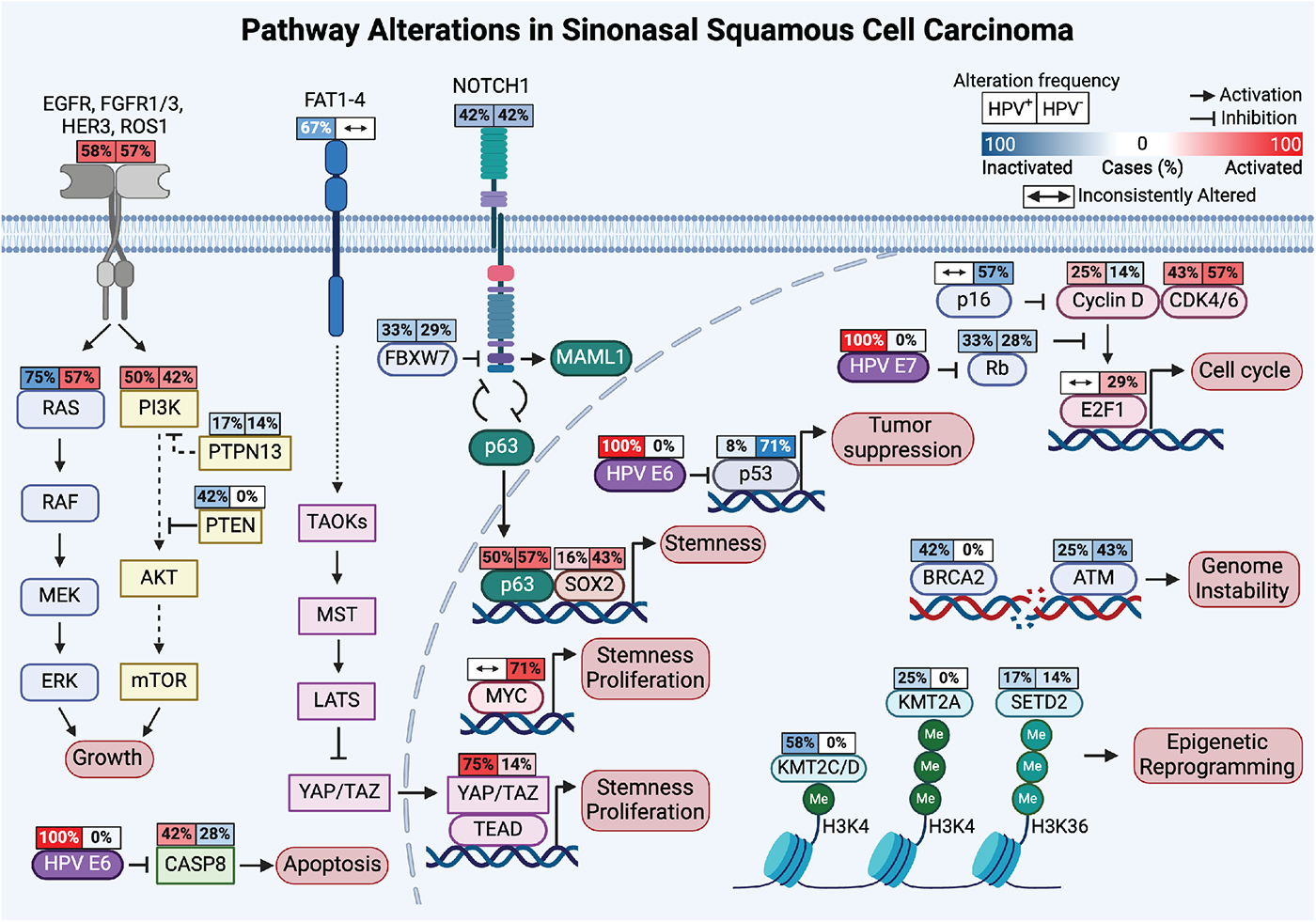
Pathway alterations in sinonasal squamous cell carcinoma. Incorporation of mutation and copy number alteration analysis reveals targetable pathways in HPV-associated SNSCC (left) and HPV-independent SNSCC (right).

We generated, to our knowledge, the first HPV-associated SNSCC cell line from patient #197, termed NCI-197 cells. This cell line was validated through both STR analysis as well as immunofluorescence staining for key markers of SNSCC (**Supplementary Figure 9**). Next, we validated the presence of active HPV16 as seen in the patient’s primary tumor using RT-PCR (data not shown). This patient’s tumor and cell line demonstrated the *PIK3CA* E542K hotspot mutation. We assessed whether small molecule inhibition of PI3K using the small molecular inhibitor alpelisib may inhibit HPV-associated SNSCC cell proliferation. Indeed, NCI-197 cells were sensitive to alpelisib as assessed by both cell viability and cell impendence assays (**Figure 7A-B**).

**Figure 7.**
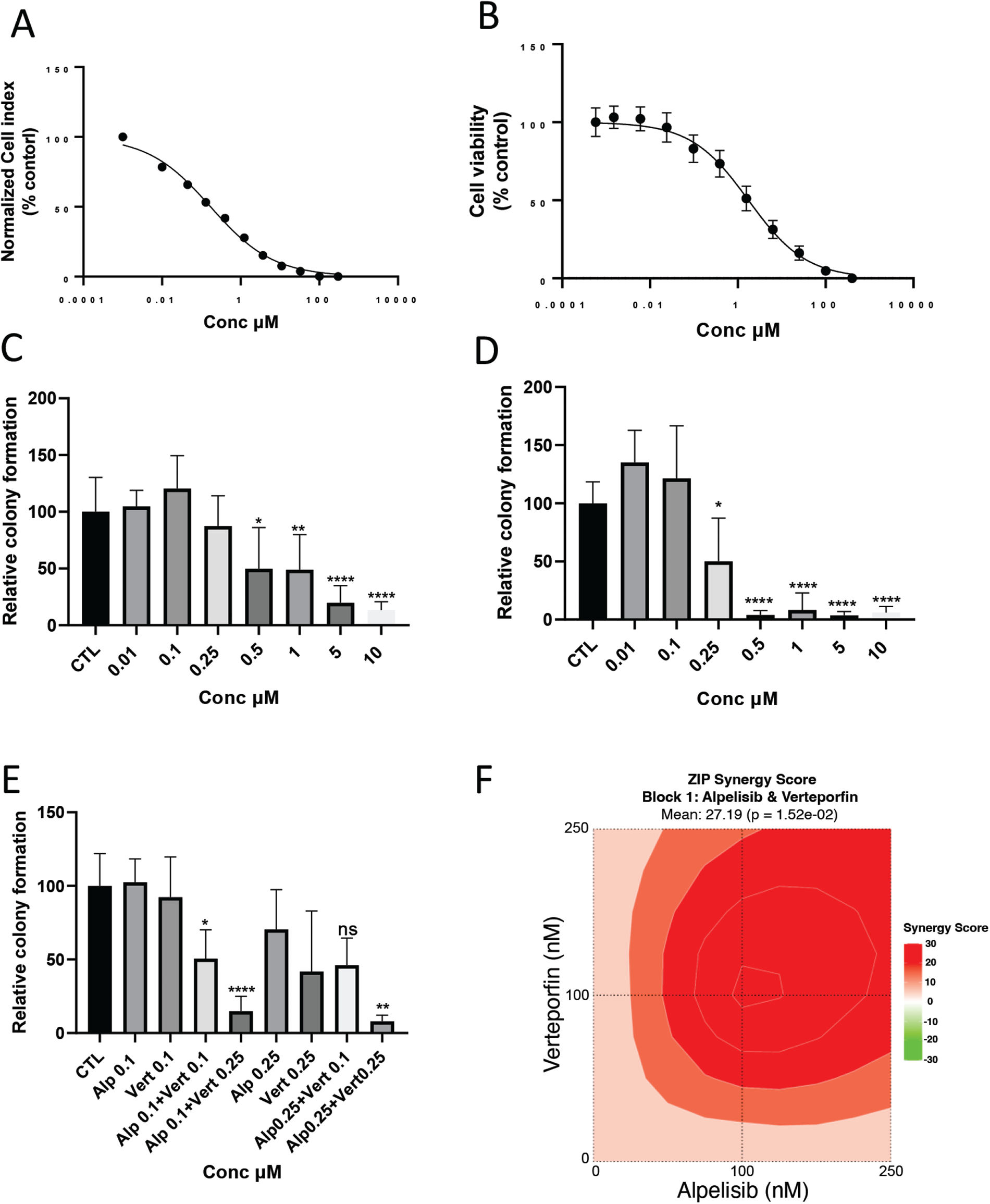
Combinatorial blockade of PI3K and YAP/TAZ synergistically inhibits HPV-associated SNSCC clonogenicity in vitro. **(A)** Impedance assay and **(B)** Cell viability analysis of NCI-197 cells 72 hours after alpelisib treatment. Data is representative of three independent experiments done in triplicates. **(C)** Dose response of alpelisib. **(D)** Dose response of verteporfin. **(E)** Combination effects of alpelisib and verteporfin assessed by colony formation assay in NCI-197 cells. Data is mean ±SEM of two independent experiments. NS = not significant; * p<0.05; ** p<0.01; **** p<0.0001. **(F)** Bliss synergy scores calculated using SyngeryFinder software for alpelisib and verteporfin dose responses (100, 250 and 500 nM).

Based on mutational and CNV analyses in HPV-associated SNSCC (**Figure 6**) we hypothesized that combinatorial inhibition of PI3K and YAP/TAZ (verteporfin) may be particularly beneficial in targeting NCI-197 cells. YAP/TAZ are known to be important in stemness and tumor initiation, thus we assessed the clonogencity of NCI-197 cells treated with small molecules inhibitors of the PI3K and TEAD (which disrupt essential YAP/TAZ-TEAD protein interactions). Interestingly, while both alpelisib and verteporfin significantly reduced colony formation individually, combinatorial treatment at lower doses significantly reduced HPV-associated SNSCC colony formation *in vitro* (**Figure 7C-E**). As assessed by SynergyFinder the combinatorial effect was determined to be a strong synergistic interaction (**Figure 7F**).

## DISCUSSION

This is the first large genome-wide analysis of HPV-associated and HPV-independent SNSCC. Prior to this study, it has been unclear whether HPV is driving tumorigenesis in SNSCC or acting as a bystander or contaminant in sinonasal malignancies(*1, 11*). We identify multiple key molecular features of HPV-driven HNSCC and CESC, including the absence of *TP53* mutations in p16 positive HPV-associated SNSCC, the presence of hot spot mutations in *PIK3CA* and *FGFR3*, APOBEC enrichment, HPV viral integration at known hotspot locations, and frequent mutations in epigenetic regulators(*6, 19, 24, 27*). These common features strongly suggest that HPV is driving HPV-associated SNSCC tumor biology rather than acting as a neutral bystander. Integration of the mutational and CNV analyses revealed combinatorial inhibition of PI3K and YAP/TAZ may be effective in reducing HPV-associated SNSCC cell clonogenicity and colony formation which was supported by synergistic small molecule inhibition of these pathways with a new HPV-associated SNSCC cell line *in vitro* (**Figure 7**).

HPV-associated SNSCC share some features of both HPV-associated HNSCC and cervical cancer. As seen for HPV-associated HNSCC and CESC, hotspot mutations were frequently identified in HPV-associated SNSCC in *PIK3CA* and *FGFR3*. Interestingly, this study identified additional recurrent hotspot mutations in multiple additional genes in HPV-associated SNSCC. Recurrent in-frame deletions were observed in *UBXN11* (c.1464_1541) in HPV-associated SNSCC and may represent an additional potential mechanism of targeting *TP53* function. A UBXD family member, *UBXN2A*, has been found through a SEP domain to bind mortalin-2(*29*). This binding results in unsequestration of p53, thus enhancing p53 activity. *UBXN11* is one of the few other UBXD family members which retains the SEP domain(*29*). Therefore, mutations in *UBXN11* resulting in decreased activity or inappropriate cell localization could potentially lead to reduced p53 activity and be an additional mechanism of p53 regulation in HPV-associated SNSCC; however, this hypothesis requires validation. Although the consequences of the *AP3S1* P158L recurrent mutation in SNSCC are unknown, recent studies have linked *AP3S1* as a driver mutation in esophageal cancer as well as a potential pan-cancer oncogene through facilitation of an immunosuppressive tumor immune microenvironment(*30, 31*). *MT-ND5* and *MT-ND4* mutations have been noted to promote tumorigenesis and metastasis in other tumor types(*32–34*). Collectively this study identifies multiple hotspot mutations which suggests these may be important oncogenic mechanisms in HPV-associated SNSCC due to their recurrent nature. However, this necessitates further investigation to determine their functional significance and biologic consequences.

The viral integration analysis in this study also identifies multiple key features of HPV-associated sinonasal malignancies including commonalities with HNSCC and CESC. Integration of HPV into or near *FGFR, TP63,* and *KLF5* is commonly observed across sinonasal malignancies, HNSCC, and CESC. Furthermore, many of the integration events on autosomes occurred at known common fragile sites. Additionally, significant enrichment of integration junctions was observed in SINE, LINE, and simple repeat sequences. Collectively these findings demonstrate many shared features of HPV integration as seen in HPV-associated HNSCC and CESC.

Viral integration analysis aided in ascertaining uncommon viral serotypes as the HPV serotype could be elucidated using this data for each HPV-associated tumor. As previously mentioned, there were 3 patients which were positive on HR-HPV ISH but negative for p16 (patient #30, 32, and 43) which were found to be positive for HPV types 114, 19, and 114, respectively. Interestingly, one of these three patients had a *TP53* missense mutation and another patient had a *TP53* nonsense mutation (**Supplementary Table 4**) and their overall survival was 9 and 13 months, respectively. These three patients also did not have any of the aforementioned recurrent mutations in *PIK3CA*, *FGFR3*, *KMT2C*, *AP3S1*, *UBXN11*, or *MT-ND5*. This suggests that SNSCC that are HR-HPV ISH positive but p16 negative have a tumor biology and behave more like HPV-independent SNSCC which may be an important clinical consideration. These data also further suggest that SNSCC should be tested for both HR-HPV ISH and p16.

There have been a few reports of HPV-associated nasolacrimal SNSCC; however, this group has not been well described due in part to their rare nature(*35–37*). Our study identified 8 patients with SNSCC arising from the nasolacrimal system. Seven of eight (87.5%) were positive for HR-HPV ISH. These 7 tumors were p16 positive and HPV16 related. HPV-associated SNSCC of the nasolacrimal system had a similar demographic and mutational profile to HPV-associated SNSCC. These clinical and molecular features suggest that HPV-associated nasolacrimal SCC be included in the HPV-associated SNSCC classification.

To our knowledge this study provides the first genome-wide analysis of a cohort of HMSC. HMSC are very different from HPV-associated SNSCC in clinical behavior and the majority are HPV33-related(*13*). Interestingly, HMSC lacked mutations in many genes including *KMT2D*, *PIK3CA*, and *FGFR3* including well defined HPV-associated hotspot locations such as E542K/E545K in *PIK3CA* and S249C in *FGFR3*. HMSC also lacked recurrent *KMT2C* N729D mutations and in frame *UBXN11* c.1464_1541 deletions in HPV-associated SNSCC but patient #252 had an *AP3S1* P158L mutation and *MT-ND5* frame-shift insertion at c.891-892 and patient #10 had a frameshift insertion in *MT-ND4* and in-frame deletion in *MT-ND5*. Interestingly, HMSC had a propensity for a higher degree of viral integration compared to other sinonasal malignancies. Two of these tumors (patient #7 and #56) had integration sites near FGFR3 with the integration event in patient #56 likely resulting in a *TACC3*::*FGFR3* fusion, a known oncogenic fusion. These data suggest that rather than direct hotspot mutations in classic HPV-associated genes such as *FGFR3*, HMSC may use alternative approaches such as viral integration to perturb these pathways. However, these patients are rare and our cohort only included 8 patients, thus these findings require further validation in additional samples.

HPV-independent SNSCC has also been poorly studied outside of targeted *TP53* sequencing. The mutational signature of SNSCC-independent patterns demonstrates many consistencies with HPV-negative HNSCC (**Supplementary Figure 8**). *TP53* mutations were frequently observed but not to the same frequency as reported for HNSCC. Similarly, *TP53* mutated HPV-independent SNSCC had worse overall survival similar to prior reports for HPV-independent HNSCC (**Figure 4A**). Furthermore, integration of mutational profiles and CNV data suggests that horizontal targeting of RAS/PI3K pathways and MYC may be an effective strategy for this tumor type (**Supplementary Figure 8**).

While this study represents the first large genomics analysis of HPV-associated and HPV-independent SNSCC, this study is limited by relatively small numbers due to the rare nature of these tumors. The findings described in this study therefore require further investigation in future multi-institutional collaborative studies. Secondly, due to the rare nature of these tumors the majority of our samples were retrospective and did not have matched normal DNA available requiring comparison to a panel of normal samples. Future studies will ideally include an increased number of samples with matched normal DNA and prospectively collected clinical information.

## MATERIALS AND METHODS

### Patient Samples

This study was approved by the Institutional Review Boards at Johns Hopkins and the National Institutes of Health. Forty-eight patients were retrospectively identified at Johns Hopkins from 2008 to 2020 with sufficient formalin-fixed paraffin embedded (FFPE) tissue and a diagnosis of squamous cell carcinoma arising from the sinonasal cavity or nasolacrimal duct or HPV-related multiphenotypic carcinoma. Of these patients, 6 had neck dissection samples from which matched normal control DNA was obtained. Sixteen patients from 2020 to 2022 were prospectively enrolled and written informed consent was obtained. Snap frozen tumor tissue was obtained along with peripheral blood mononuclear cells (PBMCs) to provide matched normal control DNA. Insufficient PBMCs for matched normal DNA comparison occurred for patient #123. Samples underwent HPV RNA in situ hybridization with a cocktail of high-risk HPV serotypes as previously described(*7*). P16 staining was performed, and positivity determined as previously described(*7*). Patients with inverted papilloma-related sinonasal squamous cell carcinoma were excluded from the study.

### DNA Extraction

DNA was extracted from 20-30mg frozen tissue samples using the AllPrep DNA/RNA Mini Kit (QIAGEN, 80204). Tissue was homogenized in 600 μL Buffer RLT Plus with β-mercaptoethanol (β-ME) using the TissueRuptor II (QIAGEN, 9002755). After centrifugation, the supernatant was transferred to AllPrep DNA spin columns. The column was centrifuged at ≥8000 x g for 30 seconds. Genomic DNA was purified by adding 500 μL Buffer AW1, centrifuging, discarding the flow-through, applying 500 μL Buffer AW2, and centrifuging again. Finally, 100 μL Buffer EB was added, incubated for 5 minutes, and centrifuged for DNA elution. The DNA was stored at –20°C.

The AllPrep DNA/RNA FFPE Kit (QIAGEN, 80234) was utilized to extract DNA from the FFPE (Formalin-Fixed Paraffin-Embedded), each containing 2-3 tissue cores. First, the samples were ground in 1.5 mL Eppendorf tubes using a disposable pestle (K7495211590).

Then, de-paraffinization was performed by adding xylene, vortexing, and centrifuging to remove the supernatant without disturbing the pellet. Ethanol was used to remove residual xylene before incubating the sample until ethanol evaporation. The pellet was then resuspended in 150 µl Buffer PKD and 10 µl proteinase K, followed by incubation at 56°C for 15 minutes and then on ice for 3 minutes. After centrifugation for 15 minutes at 20,000 x g, the supernatant was carefully transferred to a new tube, while the pellet was retained for DNA purification. For DNA extraction, the pellet was resuspended in 180 µl Buffer ATL and 40 µl proteinase K, vortexed, and incubated at 56°C for 1 hour, followed by incubation at 90°C for 2 hours without agitation. After a brief centrifugation, 200 µl Buffer AL and 200 µl ethanol (96–100%) were added to the sample, thoroughly vortexed, and transferred to a QIAamp MinElute spin column. The sample was then centrifuged for 1 minute at ≥8000 x g. Subsequent washing steps with Buffer AW1, Buffer AW2, and ethanol were performed with centrifugations at ≥8000 x g for 15 seconds each. The spin column was centrifuged at full speed for 5 minutes to dry the membrane. DNA elution was achieved by adding 40 µl Buffer ATE onto the spin column membrane, incubating for 5 minutes at room temperature, and centrifuging at full speed for 1 minute. The extracted DNA was stored at –20°C.

PBMCs were prepared for DNA extraction using the QIAamp DNA Mini and Blood Mini (QIAGEN, 51104). The PBMC sample was mixed with QIAGEN Protease and Buffer AL and then incubated at 56°C to release DNA. Ethanol was added to the lysate to promote DNA binding. The mixture was applied to QIAamp Mini spin columns and centrifuged to bind DNA to the membrane. The columns were washed with Buffer AW1 and AW2 to purify the DNA. Finally, DNA was eluted with Buffer AE and stored at -30 to -15°C.

### Whole Exome Sequencing and Whole Genome Sequencing

HPV-associated samples underwent whole genome sequencing (WGS, n=45), while HPV-independent samples underwent whole exome sequencing (WES, n=19). Libraries for WES were constructed using the Agilent SureSelect V5 post-capture beginning with a minimum of 200ng of genomic DNA and the SureSelect^XT^ Target Enrichment System for Illumina paired-end multiplexed sequencing. Libraries for WGS were constructed using TruSeq Nano DNA Library Prep kit that requires minimum 100ng of genomic DNA per sample. Paired-end sequencing was performed using the NovaSeq 6000 S4 Flow Cell with 2 × 150 cycle chemistry with an output of approximately ∼8Gb (150X) per sample for WES or ∼110Gb (30X) per sample for WGS.

### Bioinformatics Analyses

Sequencing data were demultiplexed using Illumina bcl2fastq2 (v.2.17.1.14) with default filters. Trimgalore (v0.6.7) was used to trim off adaptor sequences and low-quality bases. Reads were then aligned against the hg38 genome using BWA-MEM (v0.7.17, Sentieon 202010.02 release). Duplicate reads were removed using Picard tools (v2.9.0, Sentieon 202010.02 release). Final recalibrated alignment files were created using Genome Analysis Toolkit (GATK, v3.8.0, Sentieon 202010.02 release). To determine coverage at different levels of partitioning and aggregation, Samtools depth v1.10 & GATK DepthOfCoverage v3.8.0 were used for WES data, while Bedtools genomecov v2.30.0 was used for WGS data. Somatic variants between the tumor-normal pairs were called using GATK MuTect2 (v3.8.0, Sentieon 202010.02 release). For samples without matched normal, the panel of normal genomes from the 1000 Genomes database provided by GATK was used. GATK HaplotypeCaller (v3.8.0, Sentieon 202010.02 release) was used to call germline variants in each sample. Passed somatic and germline variants were converted into Mutation Annotation Format files using vcf2maf (v1.6.19), and then summarized and visualized using maftools (v2.10.05). Copy number analyses were performed using CNVkit (v0.9.4) for samples with matched normal, or Control-FREEC (v11.6) for samples without matched normal. Mutational signature analysis was performed using maftools (v2.18.0). The relative signature contribution barplot and boxplots were created using ggplot (v2_3.5.0). The mean comparison p-values were added to the boxplots using ggpubr (v0.6.0.999).

For mutation frequency analyses, The Cancer Genome Atlas (TCGA) HNSCC(*38*) and CESC(*25*) datasets were extracted using cBioportal(*39, 40*). HNSCC tumor HPV status was based on RNAseq reads aligning to the HPV genome, yielding 449 HPV-negative and 78 HPV-positive HNSCC tumors of all anatomic sites(*41, 42*). For CESC tumors, only cervical squamous cell carcinoma (n=141) were included. Cervical adenocarcinoma and cervical adenosquamous carcinoma as well as HPV-negative cervical tumors were excluded(*25*). Cancer-associated signaling pathways were prioritized based on pathway constituent protein products of genes frequently altered in SNSCC, HNSCC, or CESC(*25, 38, 43, 44*). Genomic alteration frequency was defined as the proportion of samples with either somatic coding sequence alteration (single nucleotide alteration or short insertion/deletion) or copy number alteration (log_2_ copy number gain/loss >0.3). Pathway figures were drawn using BioRender (biorender.com).

### HPV Integration Analysis

HPV integration analysis was conducted by first extracting the reads not mapping to the human genome or those with one unmatched mate-pair read. These reads were adapter and quality trimmed using fastp (0.23.2) remapped to a reference genome containing hg38 and all HPV genomes in the Papillomavirus Episteme (PaVE) as of 2018. From these alignments, the methods used in oncovirus_tools (https://github.com/gstarrett/oncovirus_tools, https://doi.org/10.5281/zenodo.3661416) were modified to determine integration sites for all detected HPV genomes. Integration sites were annotated using bedtools and the coordinates for all hg38 genes, fragile sites (https://webs.iiitd.edu.in/raghava/humcfs/), and repeatMasker elements.

### Establishment of HPV-associated SNSCC cell line

HPV-associated SNSCC tumor tissue was collected in PBS during surgical resection (patient #197). Tumor tissue was cleaned and cut into small fragments with a fine scissors and a single cell suspension using human tumor dissociation kit was performed per manufacturer instructions (Miltenyi biotech., 130-095-929). After tumor dissociation, RBC was lysed with ACK lysing buffer and the cells were passed through 70µM strainer. The resulting cell pellet was re-suspended in 70% ice cold basement membrane extract (Culturex reduced growth factor BME type 2, Bio-techne, 3533-010-02) in organoid growth media. Droplet of 50µl were dispensed on the bottom of pre heated 24 well plates. After incubating the plate at 37^0^C for about 20 -30 mins, 500 µl of pre-warmed patient derived organoid media was added and returned to the incubator.

Patient-derived organoid media consisted of 50% L-WRN conditioned media (obtained from ATCC and prepared as per instructions, CRL-3276) 50% advanced DMEM/F12 supplemented with 1X-B27 (life technologies, 17504-044), 1X pencillin streptomycin (life technologies), 1.25mM N-acetyl-L-Cysteine (Sigma Aldrich), 10mM nicotinamide (sigma Aldrich), 0.5 µM A83-01 (Selleckchem), 1uM SB202190, 1µM prostaglandin E2 (Selleckchem), 1µM Forskolin (Selleckchem), 0.3 µM CHIR99021 (Sigma Aldrich), 50ng/ml hEGF, 10ng/ml FGF10 and 5ng/ml FGF2 (Peprotech). After culturing organoids in BME for several passages, the organoids attached to the bottom of the plate were subsequently cultured and expanded as a 2D cell line.

The resulting patient-derived cell line was characterized by immunofluorescence staining of P63 (SantaCruz, Sc-25268), P40 and cytokeratin AE1/AE3 (SantaCruz, Sc-81714) and STR analysis was performed at Johns Hopkins Genetic Resources Core Facility. The generated cell line revives after cryopreservation and were only used for experiments up to 20 passages.

For cell viability assays, 10,000 cells were seeded in 96 well plates and different doses of alpelisib were added following overnight incubation. After 72 hours, cell viability was assessed using CellTiter-Glo 2.0 (Promega, G9241) per manufacturer instructions. Data was normalized to (100%) vehicle control and baseline (0%) 0.1% Triton X-100 and kill curves produced by fitting lines with log inhibitor versus normalized response-variable slope using GraphPad Prism software. Realtime impedance assay was performed as previously described(*45*). For colony formation clonogenicity assays, 5,000 cells were seeded in 12 well plates, allowed to attach overnight and then cells were incubated with different doses of alpelisib (Selleckchem) or verteporfin (Selleckchem) separately and in combination for 72 hours. After 72 hours, media was changed, and cells were grown for a week and stained with crystal violet. Plates were scanned and the colonies were counted.

### Statistical Analysis

For comparisons between two groups an unpaired, two-tailed, t-test was used. Log-rank (Mantel-Cox) tests were used to determine statistical significance for survival analyses. A p value significance threshold of *P*<0.05. Synergy scores were calculated using SynergyFinder (Netphar, University of Helsinki, Helsinki, Finland) with a synergy score greater than 10 indicating a strong synergistic interaction. Graphs were prepared using GraphPad Prism version 10.1.1.

## Data Availability

Sequencing data generated in this study for patients with informed consent allowing public sharing are publicly available in dbGaP. Remaining sequencing data are available upon request from the corresponding author with a completed data transfer agreement.

## Acknowledgements

The authors would like to acknowledge and thank the Johns Hopkins Experimental and Computational Genomics Core for sequencing and bioinformatics expertise. This research was supported (in part) by the Intramural Research Program of the NIH, Center for Cancer Research, National Cancer Institute. This study was supported in part by a research grant from Investigator-Initiated Studies Program of Merck Sharp & Dohme, LLC (N. London). The opinions expressed in this paper are those of the authors and do not necessarily represent those of Merck Sharp & Dohme, LLC. All other authors declare no competing interests. This study was presented at the European Skull Base Society Meeting in Maastricht, Netherlands on June 6, 2024.

