## Supplemental Figures for "Molecular patterns and mechanisms of tumorigenesis in HPV-associated and HPV-independent sinonasal squamous cell carcinoma"

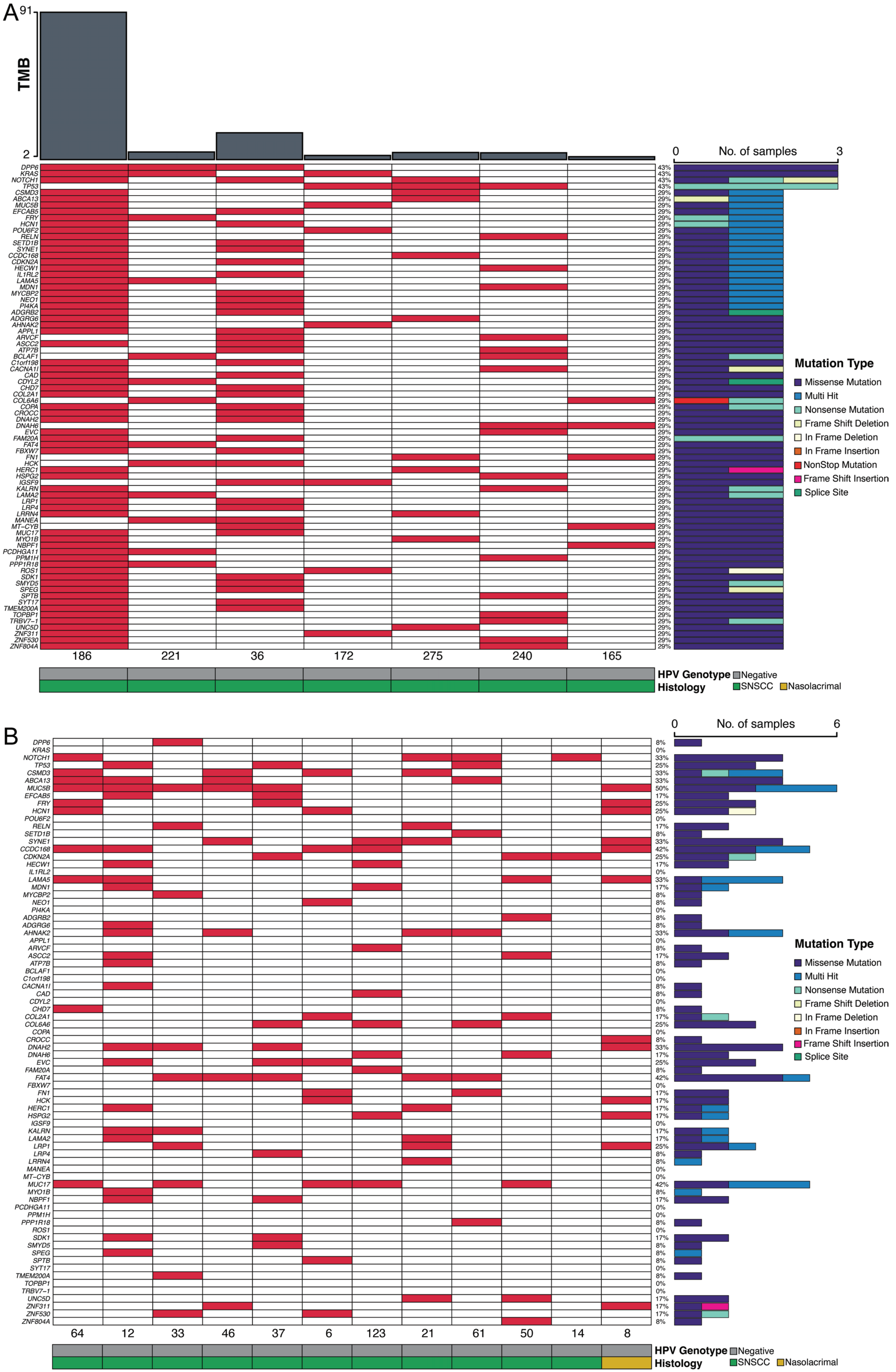

A
TMB
91
2
No. of samples
0
3
DPP6
KRAS
NOTCH1
TP53
CSMD3
ABCA13
MUC5B
EFCAB5
FRY
HCN1
POU6F2
RELN
SETD1B
SYNE1
CCDC168
CDKN2A
HECW1
IL1RL2
LAMA5
MDN1
MYCBP2
NEO1
PI4KA
ADGRB2
ADGRG6
AHNAK2
APPL1
ARVCF
ASCC2
ATP7B
BCLAF1
C1orf198
CACNA1I
CAD
CDYL2
CHD7
COL2A1
COL6A6
COPA
CROCC
DNAH2
DNAH6
EVC
FAM20A
FAT4
FBXW7
FN1
HCK
HERC1
HSPG2
IGSF9
KALRN
LAMA2
LRP1
LRP4
LRRN4
MANEA
MT−CYB
MUC17
MYO1B
NBPF1
PCDHGA11
PPM1H
PPP1R18
ROS1
SDK1
SMYD5
SPEG
SPTB
SYT17
TMEM200A
TOPBP1
TRBV7−1
UNC5D
ZNF311
ZNF530
ZNF804A
186
221
36
172
275
240
165
Mutation Type
Missense Mutation
Multi Hit
Nonsense Mutation
Frame Shift Deletion
In Frame Deletion
In Frame Insertion
NonStop Mutation
Frame Shift Insertion
Splice Site
HPV Genotype
Negative
Histology
SNSCC
Nasolacrimal
B
No. of samples
0
6
64
12
33
46
37
6
123
21
61
50
14
8
Mutation Type
Missense Mutation
Multi Hit
Nonsense Mutation
Frame Shift Deletion
In Frame Deletion
In Frame Insertion
Frame Shift Insertion
Splice Site
HPV Genotype
Negative
Histology
SNSCC
Nasolacrimal

**Supplementary Figure 1. High-throughput sequencing of HPV-independent SNSCC reveals distinct mutational patterns. (A)** Whole-exome sequencing was performed in HPV-independent SNSCC with matched normal DNA (n=7) and somatic variants assessed using a panel of normal genomes. Mutations in all genes are represented. **(B)** Whole-exome sequencing was performed in HPV-independent SNSCC without matched normal DNA (n=12) and somatic variants assessed using a panel of normal genomes. Mutations in all genes are represented.

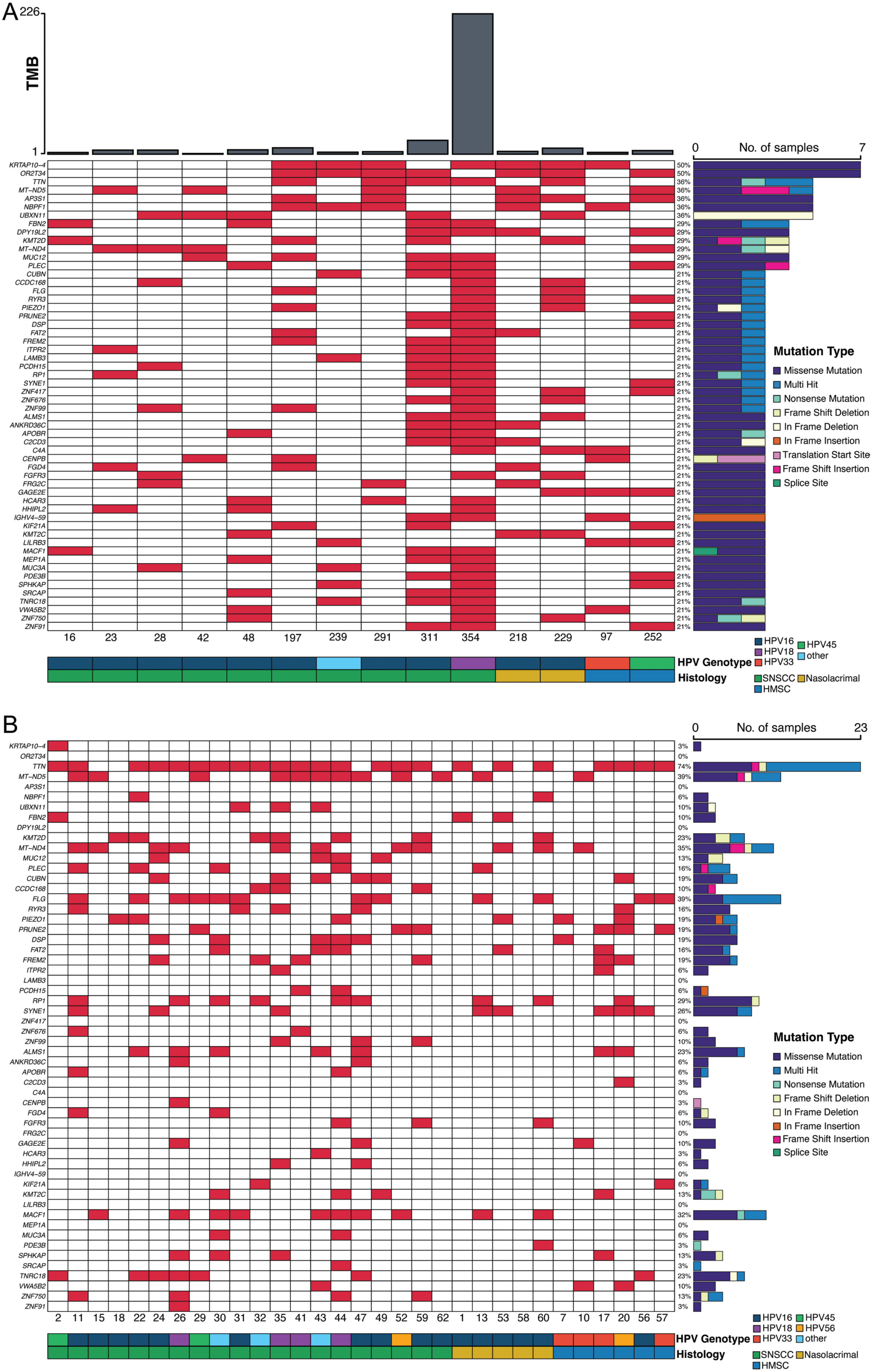
A
TMB
226
1
No. of samples
0
7
KRTAP10−4
OR2T34
TTN
MT−ND5
AP3S1
NBPF1
UBXN11
FBN2
DPY19L2
KMT2D
MT−ND4
MUC12
PLEC
CUBN
CCDC168
FLG
RYR3
PIEZO1
PRUNE2
DSP
FAT2
FREM2
ITPR2
LAMB3
PCDH15
RP1
SYNE1
ZNF417
ZNF676
ZNF99
ALMS1
ANKRD36C
APOBR
C2CD3
C4A
CENPB
FGD4
FGFR3
FRG2C
GAGE2E
HCAR3
HHIPL2
IGHV4−59
KIF21A
KMT2C
LILRB3
MACF1
MEP1A
MUC3A
PDE3B
SPHKAP
SRCAP
TNRC18
VWA5B2
ZNF750
ZNF91
16 23 28 42 48 197 239 291 311 354 218 229 97 252
Mutation Type
Missense Mutation
Multi Hit
Nonsense Mutation
Frame Shift Deletion
In Frame Deletion
In Frame Insertion
Translation Start Site
Frame Shift Insertion
Splice Site
HPV Genotype
HPV16
HPV18
HPV33
HPV45
other
Histology
SNSCC
Nasolacrimal
HMSC
B
No. of samples
0
23
2 11 15 18 22 24 26 29 30 31 32 35 41 43 44 47 49 52 59 62 1 13 53 58 60 7 10 17 20 56 57
Mutation Type
Missense Mutation
Multi Hit
Nonsense Mutation
Frame Shift Deletion
In Frame Deletion
In Frame Insertion
Frame Shift Insertion
Splice Site
HPV Genotype
HPV16
HPV18
HPV33
HPV45
HPV56
other
Histology
SNSCC
Nasolacrimal
HMSC

**Supplementary Figure 2. High-throughput sequencing of HPV-associated SNSCC reveals distinct mutational patterns. (A)** Whole-genome sequencing was performed in HPV-associated SNSCC with matched normal DNA (n=12) and HMSC (n=2) and somatic variants were assessed. Mutations in all genes are represented. **(B)** Whole-genome sequencing was performed in HPV-associated SNSCC without matched normal DNA (n=25) and HMSC (n=6) and somatic variants assessed using a panel of normal genomes. Mutations in all genes are represented.

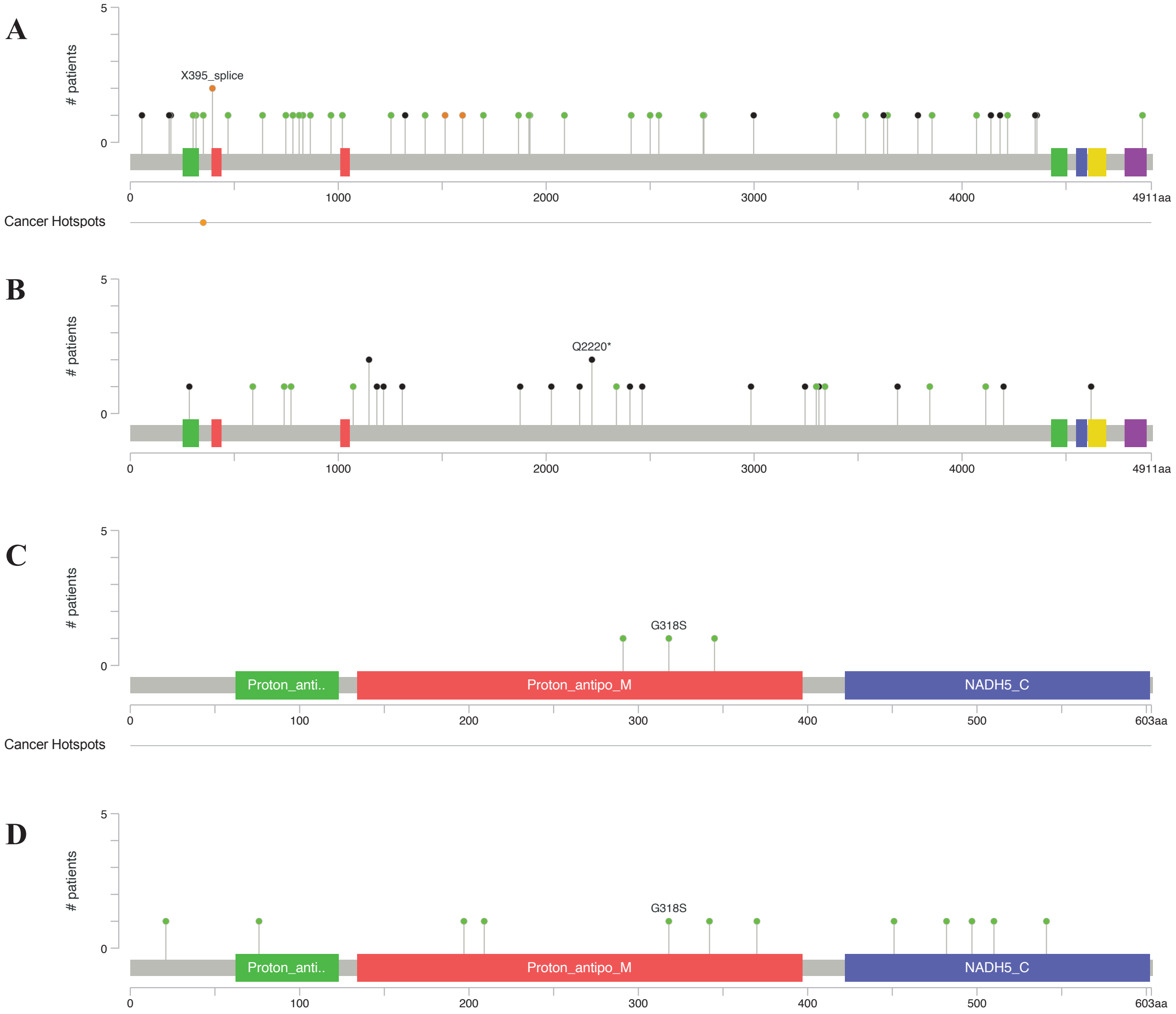


**Supplementary Figure 3. HNSCC and CSCC lack *KMT2C* N729D mutations and *MT-ND5* frameshift insertion/deletions at c.891-892 or c.1278-1279. (A)** TCGA was evaluated for *KMT2C* mutations in HNSCC and 0/46 N729D mutations were observed. **(B)** TCGA was evaluated for *KMT2C* mutations in CSCC and 0/37 N729 mutations were observed. **(C)** TCGA was evaluated for *MT-ND5* mutations in HNSCC and 0/3 *MT-ND5* c.891-892 or c.1278-1279 frameshift insertion/deletions were noted. **(D)** TCGA was evaluated for *MT-ND5* mutations in CSCC and 0/12 *MT-ND5* c.891-892 or c.1278-1279 frameshift insertion/deletions were noted.

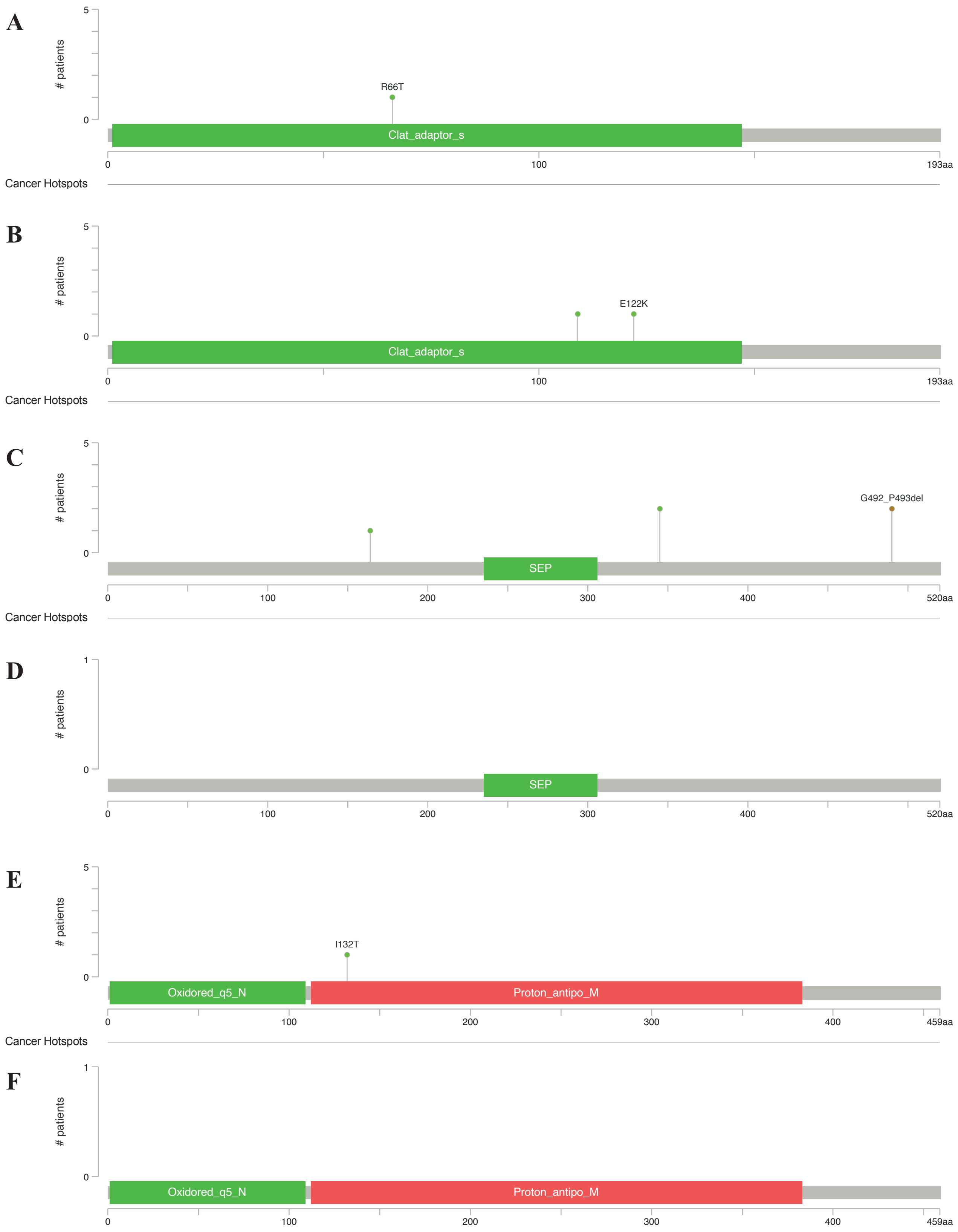


**Supplementary Figure 4. HNSCC and CSCC either lack or have few mutations in *AP3S1*, *UBXN11*, and *MT-ND4*.** TCGA was evaluated for *AP3S1* mutations in HNSCC **(A)** and CSCC **(B)** and few mutations were observed. TCGA was evaluated for *UBXN11* mutations in HNSCC **(C)** and CSCC **(D)** and few mutations were observed. TCGA was evaluated for *MT-ND4* mutations in HNSCC **(E)** and CSCC **(F)** and few mutations were observed. None of the recurrent changes in these genes were seen as in HPV-associated SNSCC.

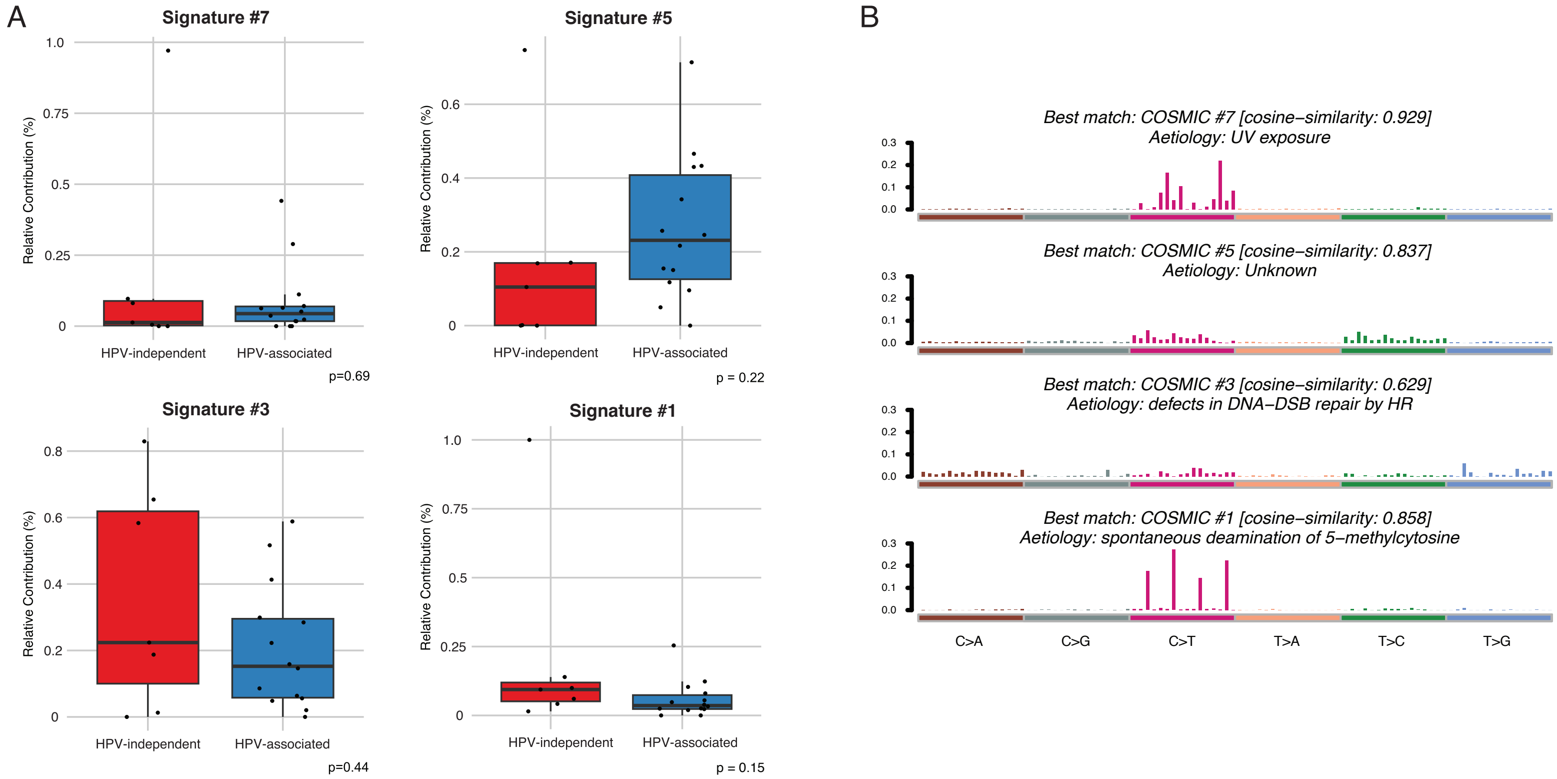


**Supplementary Figure 5. Additional pathway enrichment in HPV-independent and HPV-associated SNSCC.**
**(A)** Additional dox plot comparisons of pathway enrichment in HPV-independent and HPV-associated SNSCC.
**(B)** Signature profiles in the HPV-associated SNSCC cohort.

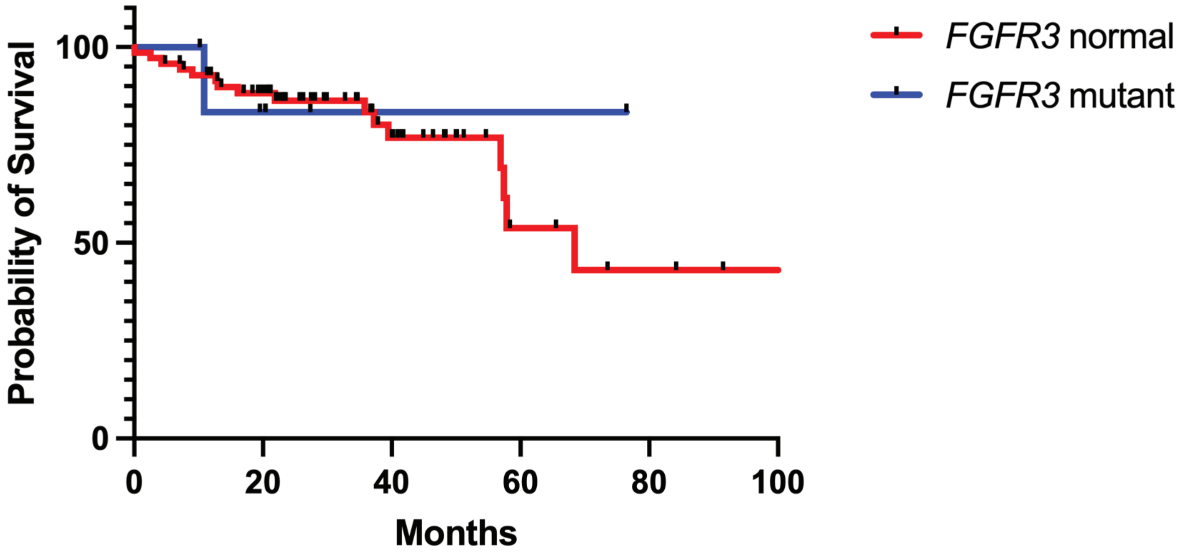


**Supplementary Figure 6. No association with overall survival and *FGFR3* mutation status in HNSCC. (A)** HPV-associated HNSCC from TCGA with *FGFR3* mutations and overall survival ($P$ = 0.718).

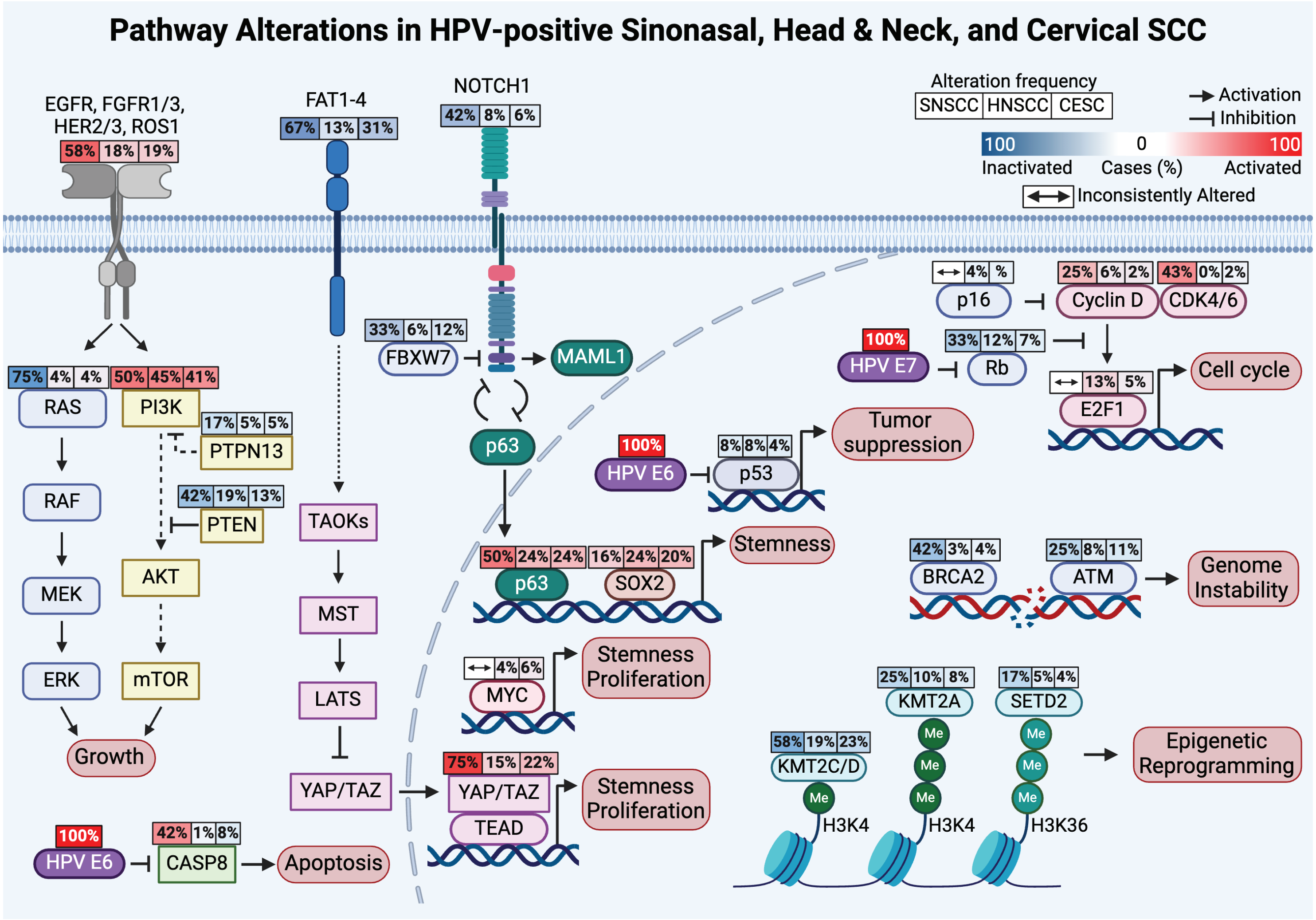


**Supplementary Figure 7. Pathway alteration comparison of HPV-associated disease.** Incorporation of mutation and copy number alteration analysis reveals targetable pathways enhanced in HPV-associated SNSCC (left), compared to HPV-associated HNSCC (middle), and HPV-associated CESC (right). Figure drawn using BioRender (biorender.com).

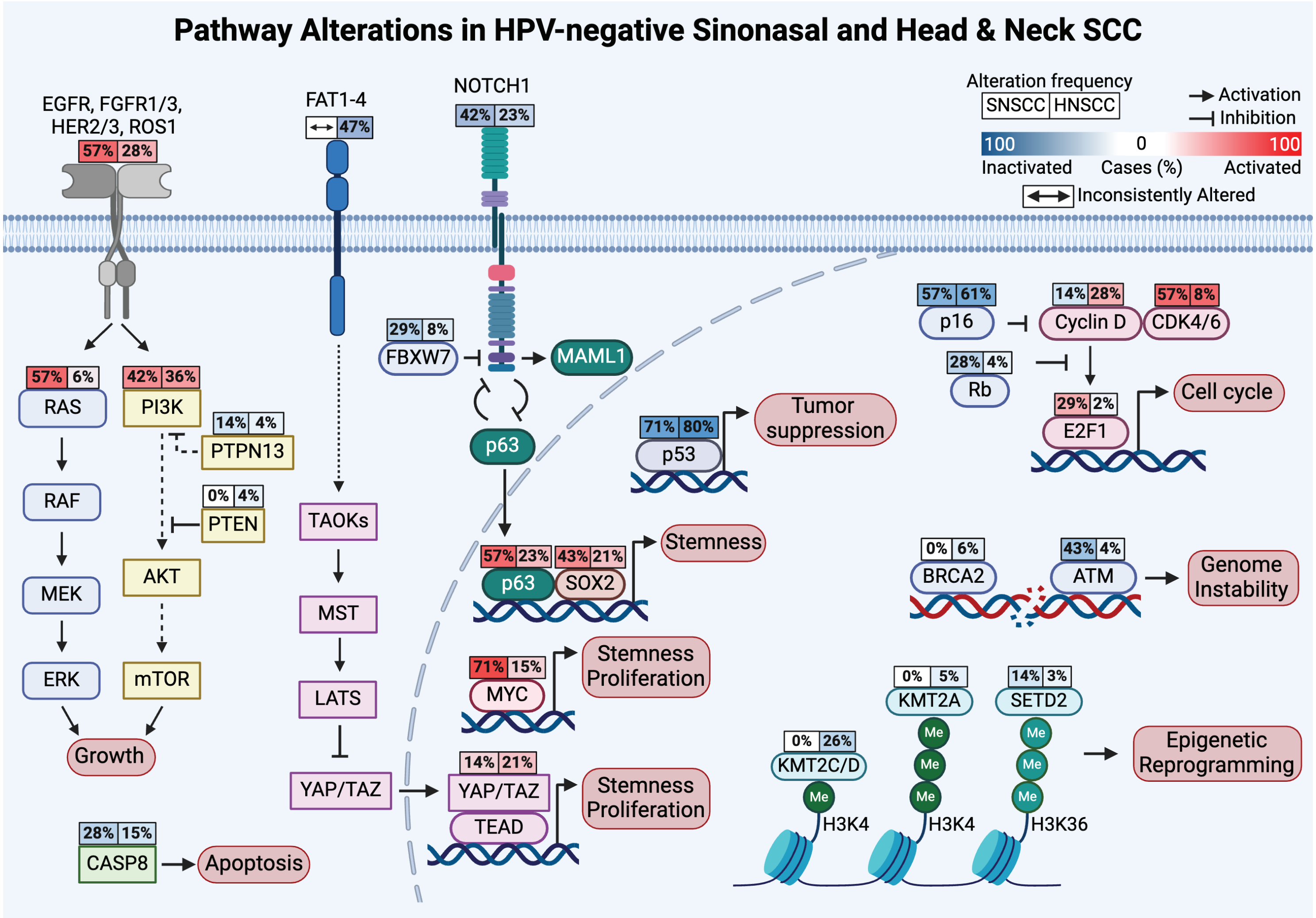


**Supplementary Figure 8. Pathway alteration comparison of HPV-independent disease.** Incorporation of mutation and copy number alteration analysis reveals targetable pathways enhanced in HPV-independent SNSCC (left) compared to HPV-independent HNSCC (right). Figure drawn using BioRender (biorender.com).

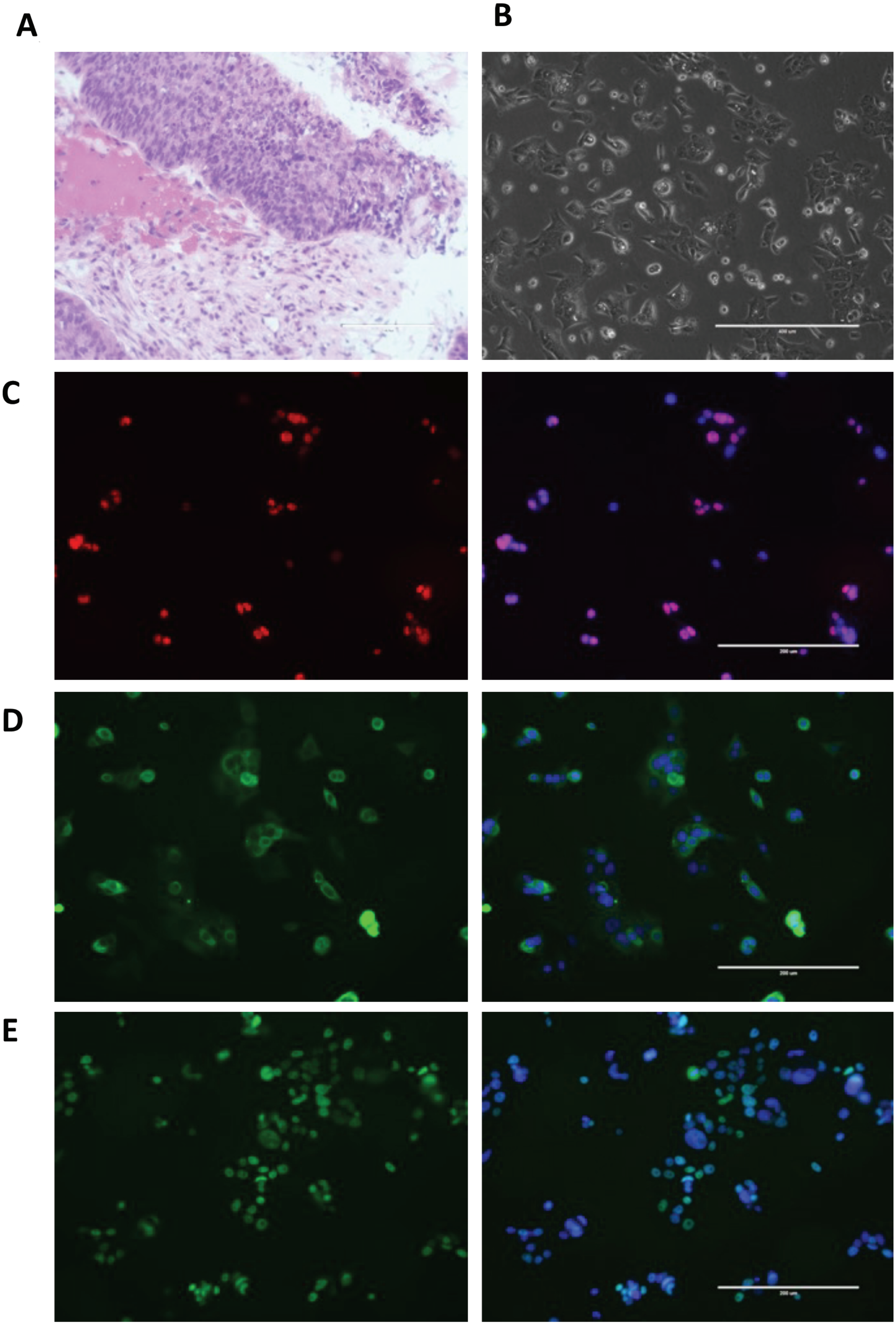


**Supplementary Figure 9. Validation of HPV-associated SNSCC cell line. (A)** H&E staining of original primary tumor. **(B)** Bright field image and Immunofluorescence staining for **(C)** P63, **(D)** cytokeratin AE1/AE3, and **(E)** P40 of cell line established from primary tumor. Scale bar in A and B are 400 μm and 200 μm in C-E.
